# Lipin-1-cardiolipin axis sustains mitochondrial respiration and efferocytosis in macrophages

**DOI:** 10.64898/2026.11.23.768679

**Authors:** Oluwakemi O. Igiehon, Cora R. Palmer, Javier Nion-Fieira, Temitayo T. Bamgbose, Zachariah Martin, Robert M. Schilke, Chowdury S. Abdullah, Lindsey Key, Andrii Andriichuk, Rajan Pandit, Kevin J. McCarthy, Deborah McCarthy, Md Shenuarin Bhuiyan, Arif Yurdagul, Matthew D. Woolard

## Abstract

Defective macrophage clearance of apoptotic cells contributes to persistent inflammation and cardiometabolic disease. Efferocytosis requires mitochondrial oxidative metabolism, but how macrophages maintain respiratory capacity during resolution remains unclear. Here, we show that lipin-1 supports efferocytosis through a cardiolipin-associated mitochondrial program. In IL-4-stimulated macrophages, lipin-1 deficiency reduced cardiolipin abundance and selectively lowered the abundance of inner mitochondrial membrane proteins, including respiratory-chain components, consistent with decreased mitochondrial respiratory capacity. These defects occurred without broad suppression of canonical metabolic transcriptional programs, and experimentally altering mitochondrial morphology failed to reproduce or rescue the functional phenotype. Disrupting cardiolipin synthesis impaired efferocytosis, whereas the cardiolipin-interacting peptide SS-31 improved mitochondrial respiration and apoptotic-cell clearance in lipin-1-deficient macrophages. Systemic SS-31 treatment also reduced apoptotic-cell accumulation after dexamethasone-induced thymic injury. Together, these findings implicate cardiolipin homeostasis in lipin-1-dependent macrophage bioenergetics and efferocytosis, identifying a potentially actionable pathway while leaving the mechanism connecting lipin-1 to cardiolipin abundance unresolved.

**Graphical Abstract:** 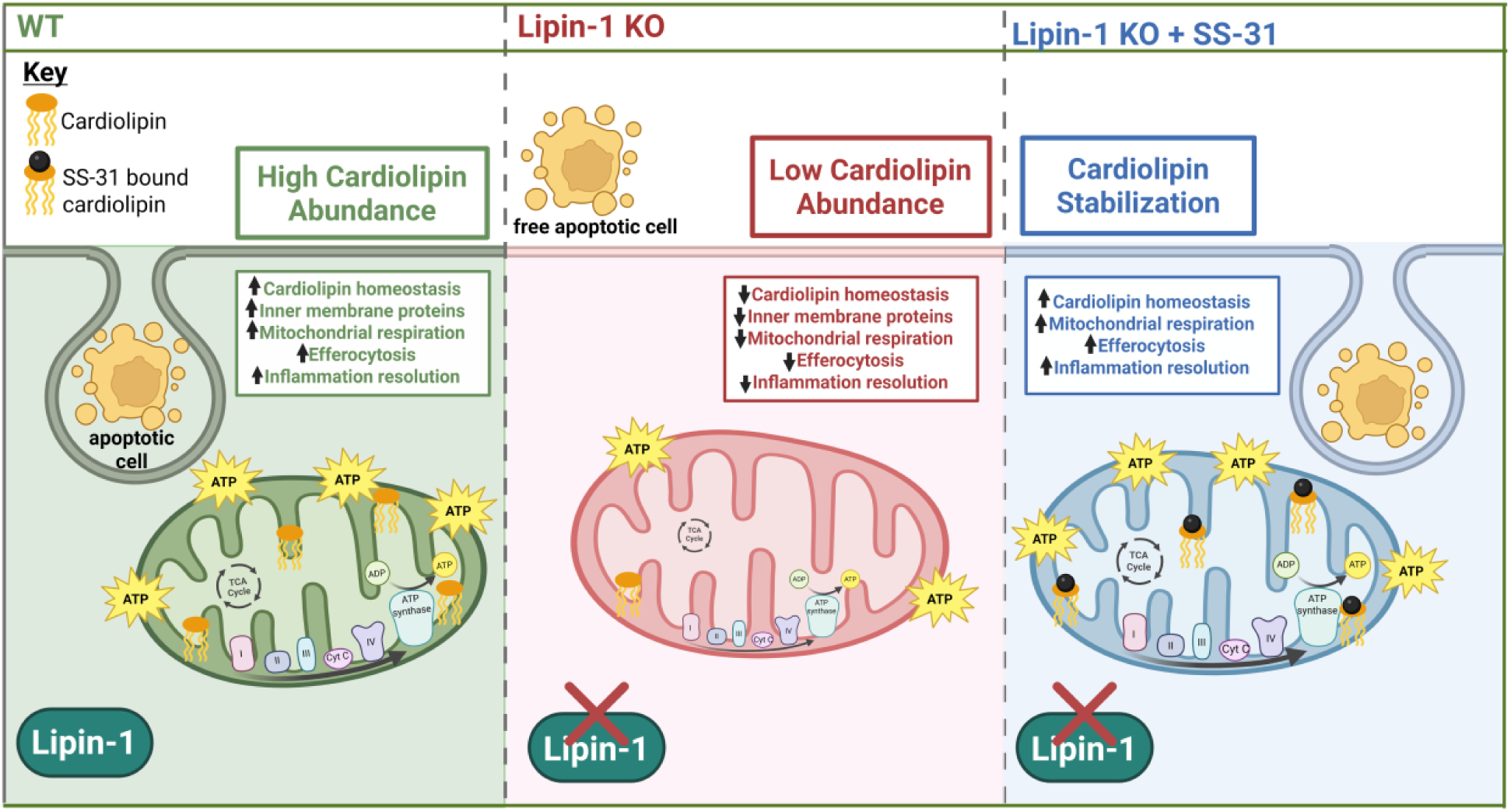

## INTRODUCTION

Failure to effectively resolve inflammation contributes to chronic inflammation associated with atherosclerotic cardiovascular disease, metabolic dysfunction-associated steatotic liver disease (MASLD), insulin resistance, diet-induced obesity, and several forms of cancer ^1,2^. Macrophages play central roles in initiating and resolving inflammation owing to their functional plasticity. During the resolution phase, pro-resolving macrophages clear apoptotic cells through efferocytosis, secrete anti-inflammatory cytokines such as IL-10 and TGF-β, and generate specialized pro-resolving mediators such as resolvins, protectins, and maresins, thereby promoting the restoration of tissue homeostasis ^3–6^. These functions are supported by extensive metabolic remodeling. In particular, mitochondrial fatty acid β-oxidation and oxidative phosphorylation (OXPHOS) support efferocytosis and other pro-resolving responses ^3,7,8^. Although mitochondrial metabolism has emerged as a key determinant of macrophage efferocytosis and inflammation resolution, the mechanisms that maintain mitochondrial oxidative capacity during these responses remain incompletely defined.

Lipin-1, a phosphatidic acid phosphatase and transcriptional co-regulator, is well positioned to regulate macrophage mitochondrial metabolism because it integrates cellular lipid metabolism with mitochondrial bioenergetics. We previously demonstrated that macrophage-associated lipin-1 promotes efferocytosis and supports inflammation resolution and atheroprotection ^8,9^. Furthermore, we showed that macrophage lipin-1 restrains ceramide accumulation while supporting fatty acid β-oxidation and mitochondrial OXPHOS, metabolic programs that support IL-4-mediated pro-resolving macrophage responses, including efferocytosis ^7,8^. Collectively, these findings identify lipin-1 as an important regulator of macrophage lipid metabolism and mitochondrial function, yet how lipin-1 maintains mitochondrial homeostasis to support macrophages and inflammation resolution remains unclear.

Cardiolipin (CL) is a tetra-acylated diphosphatidylglycerol phospholipid and constitutes 15-20% of total mitochondrial phospholipid content and is highly enriched in the inner mitochondrial membrane ^10^. Cardiolipin supports inner mitochondrial membrane architecture, facilitates cristae formation, and promotes the assembly and stabilization of electron transport chain (ETC) supercomplexes, thereby supporting efficient OXPHOS and ATP production ^11^. Consequently, disruption of cardiolipin synthesis, remodeling, or maintenance impairs ETC function and compromises OXPHOS ^12,13^. Cardiolipin exerts context-dependent effects on inflammatory signaling that are influenced by its abundance and acyl-chain composition. Exogenous unsaturated cardiolipin attenuates TLR4-mediated inflammatory responses to LPS in macrophages and can augment phagocytic activity in murine microglia and macrophages, including PMA-differentiated THP-1 cells ^14–17^. Endogenous mitochondrial cardiolipin can also support metabolic reprogramming during pro-inflammatory macrophage activation and contribute to NLRP3 inflammasome activation ^18,19^. Despite these advances, the role of CL in macrophage pro-resolving responses, including efferocytosis and inflammation resolution, remains largely unexplored. Notably, cardiac lipin-1 deficiency reduces cardiolipin abundance and impairs mitochondrial respiration despite increasing phosphatidic acid (PA), a precursor of CL biosynthesis ^20^. This paradox suggests that lipin-1 may influence cardiolipin homeostasis through mechanisms beyond bulk PA availability.

Based on these observations, we hypothesized that lipin-1 maintains mitochondrial homeostasis to support mitochondrial function and macrophage efferocytosis. We therefore investigated how lipin-1 deficiency alters mitochondrial organization, inner mitochondrial membrane protein composition, and phospholipid homeostasis, as well as the functional consequences of these alterations on mitochondrial respiration and efferocytosis in response to IL-4. Our findings identify cardiolipin homeostasis as a key component of lipi-1-dependent regulation of mitochondrial function and macrophage efferocytosis.

## RESULTS

### Lipin-1-dependent regulation of macrophage lipid metabolism is not explained by broad changes in metabolic gene expression

We previously demonstrated that lipin-1, independent of its enzymatic activity, enhances IL-4-induced efferocytosis and OXPHOS, whereas loss of lipin-1 increases ceramide abundance in IL-4-stimulated macrophages ^7,8^. In hepatocytes, lipin-1 functions as a transcriptional co-regulator that interacts with PPARα and PGC-1α to amplify transcriptional programs involved in fatty acid β-oxidation and mitochondrial oxidative metabolism ^21^. Nuclear lipin-1 also suppresses SREBP-dependent transcription, thereby limiting the expression of genes involved in fatty acid and cholesterol synthesis ^22,23^. Together, these transcriptional regulatory activities could favor fatty acid utilization over lipid synthesis, potentially contributing to the metabolic remodeling that supports macrophage efferocytosis. We therefore tested whether the metabolic phenotype of lipin-1-deficient macrophages was associated with alterations in canonical PPAR-and SREBP-dependent transcriptional programs.

We performed bulk RNA sequencing of untreated and IL-4-treated (40 ng/mL for 4 hours) WT and lipin-1mKO BMDMs using 3 biological replicates per condition. Using an FDR threshold of 0.10 and a minimum absolute fold-change cutoff of 1.2, we identified 419 differentially expressed genes in IL-4-treated lipin-1mKO macrophages, including 284 downregulated and 135 upregulated genes. Contrary to our expectation, GO Biological Process overrepresentation analysis did not identify preferential enrichment of lipid or mitochondrial metabolic pathways among genes downregulated in lipin-1mKO macrophages. Instead, enriched terms included immune-response pathways, including type I interferon signaling (Figure 1A–C). Based on the known effects of lipin-1 on PPAR and SREBP transcriptional programs, we expected genes downregulated in IL-4-treated lipin-1mKO BMDMs to be PPAR target genes, and upregulated genes to be SREBP target genes. Surprisingly, PPARα and PPARγ were not among the most prominent transcription factors identified by transcription factor enrichment analysis using the Enrichr ENCODE TF ChIP-seq library. Likewise, SREBP target genes were not significantly enriched within either differentially expressed gene sets (Figure 1D, E). Instead, genes downregulated in IL-4-treated lipin-1mKO BMDMs were significantly enriched for genes associated with IRF1 and IRF8 binding in the ENCODE ChIP-seq datasets (Figure 1D).

**Figure 1.**
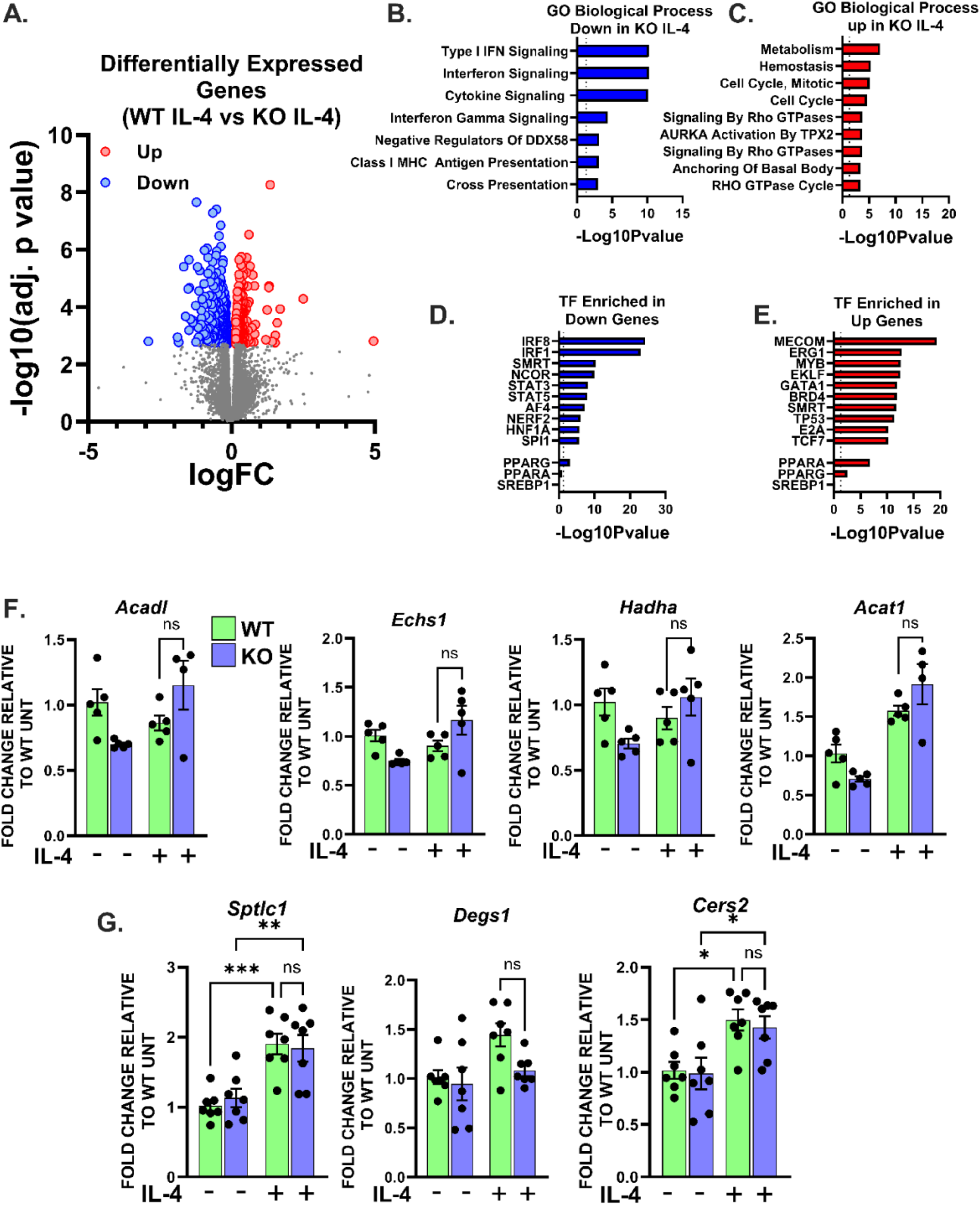
Lipin-1-dependent regulation of macrophage lipid metabolism is not explained by broad changes in metabolic gene expression. WT and lipin-1mKO bone marrow-derived macrophages (BMDMs) were analyzed following IL-4 stimulation. (A) Volcano plot of differentially expressed genes in IL-4-treated WT versus lipin-1mKO BMDMs. RNA sequencing was performed after 4 h of IL-4 treatment (40 ng/mL; n = 3 biological replicates per genotype). Differentially expressed genes were defined by a Benjamini-Hochberg-adjusted false-discovery rate (FDR) < 0.10 and an absolute fold change ≥1.2; genes not meeting these criteria are shown in gray. (B and C) Gene Ontology Biological Process overrepresentation analysis of genes downregulated (B) or upregulated (C) in IL-4-treated lipin-1mKO BMDMs relative to WT BMDMs. (D and E) Transcription-factor target enrichment analysis of genes downregulated (D) or upregulated (E) in lipin-1mKO BMDMs using Enrichr and the ENCODE TF ChIP-seq library. (F and G) qRT-PCR analysis of genes involved in mitochondrial fatty-acid β-oxidation (F; *Acadl, Echs1, Hadha,* and *Acat1*) and ceramide synthesis (G; *Sptlc1, Degs1,* and *Cers2*) in untreated or IL-4-treated WT and lipin-1mKO BMDMs. Expression was normalized to *Gapdh* and expressed relative to untreated WT BMDMs. Each point represents an independent BMDM preparation [n = 4-7 per group]. Data in (F) and (G) are mean ± SEM and were analyzed using one-way ANOVA in GraphPad Prism. \**p* < 0.05, \*\**p* < 0.01, \*\*\**p* < 0.001, \*\*\*\**p*< 0.0001; ns, not significant.

To independently assess selected metabolic pathways, we performed qRT-PCR on mitochondrial fatty-acid oxidation-associated genes (*Acadl, Echs1, Hadha,* and *Acat1*) and ceramide synthesis-associated genes (*Sptlc1, Degs1,* and *Cers2*) from new samples. IL-4 increased the expression of several ceramide-biosynthetic genes (*Sptlc1* and *Cers2)*; however, their expression did not differ significantly between WT and lipin-1mKO BMDMs (Figure 1G). Thus, the previously observed lipin-1-dependent differences in ceramide abundance are unlikely to result from altered transcription of these biosynthetic enzymes. Similarly, the expression of the selected β-oxidation genes did not differ significantly between WT and lipin-1mKO BMDMs (Figure 1F).

Together, these findings argue against disruption of canonical PPAR- and SREBP-dependent transcriptional programs as the primary mechanism underlying the altered metabolic phenotype of lipin-1-deficient macrophages. We therefore investigated whether lipin-1 regulates mitochondrial biology through alternative mechanisms that support mitochondrial function and macrophage efferocytosis.

### Lipin-1-deficient macrophages exhibit elongated mitochondrial networks

Because broad changes in metabolic gene expression did not explain the lipin-1-dependent metabolic phenotype, we next investigated whether lipin-1 deficiency alters mitochondrial morphology. Mitochondrial morphology is closely linked to mitochondrial bioenergetics and macrophage function, with DRP1-mediated mitochondrial fission supporting continued uptake of apoptotic cells during efferocytosis ^24–27^. In addition, lipin-1 overexpression has been shown to increase mitochondrial fragmentation in HeLa ^28^, suggesting that lipin-1 may influence mitochondrial dynamics. Thus, we examined mitochondrial morphology in WT and lipin-1mKO BMDMs.

We assessed mitochondrial morphology in WT and lipin-1mKO BMDMs by TOM-20 immunostaining, confocal microscopy, and quantitative image analysis (Figure 2A). IL-4-treated lipin-1mKO BMDMs exhibited significantly increased mitochondrial length compared with WT BMDMs (Figures 2B, 2D, & S1A), whereas mitochondria in WT cells had higher circularity following IL-4 treatment (Figure 2C & S1B). These findings indicate a more elongated mitochondrial network in lipin-1mKO BMDMs and are phenotypically consistent with prior observations that lipin-1 overexpression promotes mitochondrial fragmentation in non-macrophage cells. Because lipin-1 deficiency has previously been associated with impaired mitophagy and mitochondrial accumulation in glycolytic muscle, we determined whether the elongated mitochondrial morphology in lipin-1mKO BMDMs was accompanied by increased mitochondrial genome abundance. We quantified mtDNA relative to nuclear DNA. The mtDNA:nDNA ratio did not differ between WT and lipin-1-deficient BMDMs (Figure 2E), indicating that the altered mitochondrial morphology was not associated with a detectable increase in mitochondrial genome abundance. Although this finding argues against gross accumulation of mtDNA-containing mitochondrial material, it does not exclude changes in mitophagic flux, mitochondrial biogenesis, or mitochondrial turnover.

**Figure 2.**
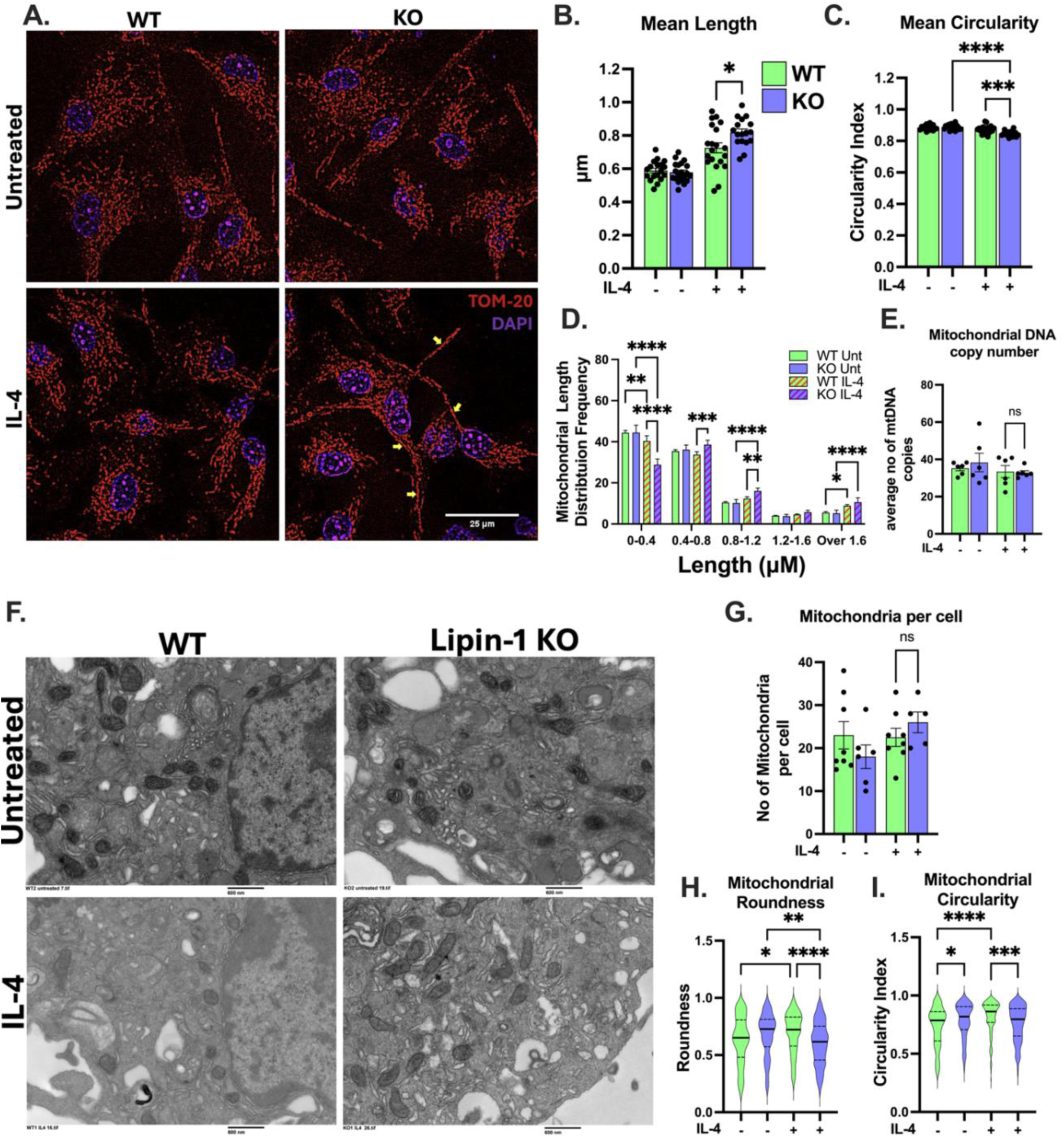
Lipin-1 deficiency is associated with an elongated mitochondrial morphology. WT and lipin-1mKO BMDMs were treated with or without IL-4 (40 ng/mL) for 6 h. (A) Representative confocal images of mitochondria identified by TOM-20 immunofluorescence (red); nuclei were stained with DAPI (blue). Scale bar, 25 μm. Yellow arrows indicate elongated mitochondria. (B and C) Quantification of mean mitochondrial length (B) and mean circularity (C) per field. (D) Distribution of mitochondrial lengths across the indicated size bins. (E) Relative mitochondrial DNA copy number determined by qPCR using two mitochondrial-to-nuclear DNA ratios and reported as the mean of the two measurements. (F) Representative transmission electron micrographs of mitochondria from untreated and IL-4-treated WT and lipin-1 KO BMDMs. Scale bar, 600 nm. (G) Number of mitochondrial profiles per complete cellular section determined from transmission electron microscopy images. (H and I) Quantification of mitochondrial roundness (H) and circularity (I) from transmission electron micrographs. (B, C, H & I) Violin plots show the distribution of mitochondrial measurements (center lines are displayed in bold, and quartiles are displayed in dashes). Data in bar graphs are shown as means ± SEM with n= 5-7 (E & G). Statistical analyses used are one-way ANOVA (B, C, G, H & I), two-way ANOVA (D), and unpaired *t* test (E). \**p* < 0.05, \*\**p* < 0.01, \*\*\**p* < 0.001, \*\*\*\**p*< 0.0001; ns, not significant.

To validate and extend the confocal microscopy findings at the ultrastructural level, we examined mitochondria by transmission electron microscopy and quantified the number of mitochondrial profiles per cellular section, as well as mitochondrial roundness and circularity (Figure 2F) ^29^. The number of mitochondrial profiles did not differ significantly between WT and lipin-1mKO BMDMs (Figure 2G). Following IL-4 treatment, mitochondrial profiles in WT BMDMs were significantly rounder and more circular than those in lipin-1mKO BMDMs (Figure 2H, I), consistent with a less elongated mitochondrial morphology and supporting the confocal microscopy findings. Mitochondrial roundness and circularity were quantified as complementary morphological metrics. Higher values for both parameters indicate a more rounded, less elongated mitochondrial shape, consistent with mitochondrial fragmentation. Conversely, lower values reflect a more elongated mitochondrial network. ^30^. Consistent with increased mitochondrial length, IL-4-treated lipin-1mKO BMDMs also exhibited significantly greater mitochondrial area and aspect ratio, indicative of altered mitochondrial architecture and enhanced elongation **(Figure S1C-E)** ^31^. Together, these findings demonstrate that lipin-1 deficiency is associated with an elongated, less circular mitochondrial morphology without a detectable increase in mtDNA copy number or mitochondrial profiles per cellular section. These observations prompted us to determine whether the altered morphology contributes functionally to the lipin-1-dependent defects in efferocytosis and oxidative phosphorylation.

### Altered mitochondrial morphology does not account for lipin-1-dependent defects in efferocytosis

Lipin-1mKO BMDMs exhibited an elongated mitochondrial network, but it remains unclear whether this morphological phenotype contributed to impaired efferocytosis. Because DRP1-mediated mitochondrial fission facilitates the continued uptake of apoptotic cells ^26^, we tested whether experimentally altering mitochondrial morphology would reproduce or rescue the lipin-1-dependent functional phenotypes. We depleted OPA1 in lipin-1mKO BMDMs to reduce mitochondrial fusion and promote fragmentation and depleted DRP1 in WT BMDMs to reduce fission and promote elongation, using 200 nM siRNA targeting each protein. We confirmed OPA-1 and DRP1 depletion by immunoblotting, and OPA-1 depletion significantly reduced mitochondrial length in lipin-1mKO BMDMs, whereas DRP1 depletion increased mitochondrial length in WT BMDMs, as shown by TOM-20 staining and immunofluorescence microscopy (Figure 3A-C). We next determined whether these reciprocal changes in mitochondrial morphology altered IL-4-induced efferocytosis. OPA1 depletion did not restore efferocytosis in lipin-1mKO BMDMs, indicating that promoting mitochondrial fragmentation through loss of OPA1 was insufficient to correct the lipin-1-dependent defect under these assay conditions. Conversely, DRP1 depletion did not significantly reduce efferocytosis in WT BMDMs (Figure 3D, E). Thus, experimentally induced mitochondrial elongation did not reproduce the lipin-1mKO phenotype, although these results do not exclude a role for DRP1-mediated fission during high-burden or serial efferocytosis.

**Figure 3.**
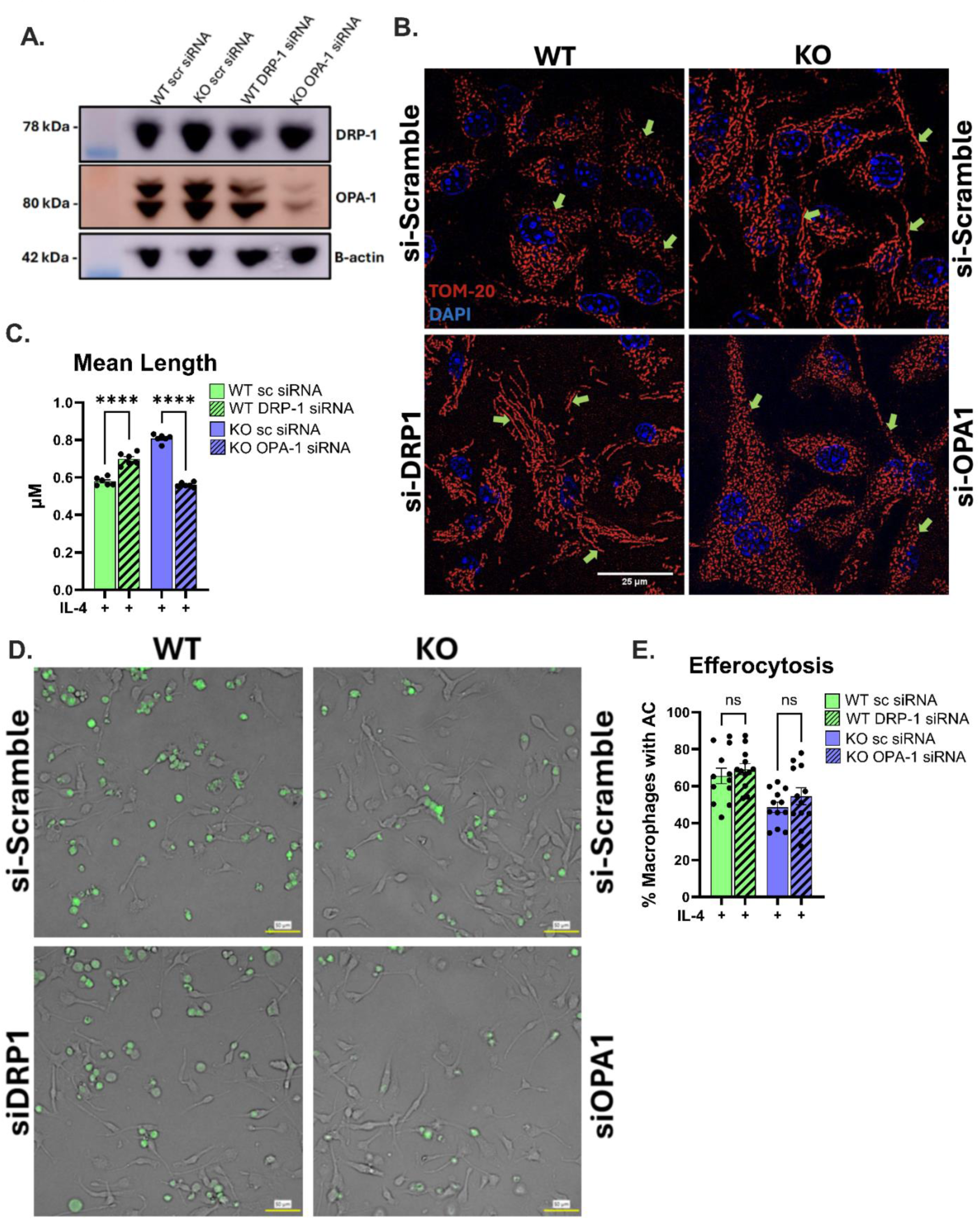
Manipulation of mitochondrial fission and fusion does not reproduce or rescue the lipin-1-dependent functional phenotype. (A) Representative immunoblots confirming siRNA-mediated depletion of DRP1 in WT BMDMs and OPA1 in lipin-1mKO BMDMs. β-Actin served as the loading control. (B) Representative TOM-20 immunofluorescence images demonstrating mitochondrial morphology following DRP1 depletion in WT BMDMs or OPA1 depletion in lipin-1mKO BMDMs. TOM-20 is shown in red and nuclei in blue. Scale bar, 25 μm. Green arrows highlight mitochondrial lengths. (C) Quantification of mean length of TOM-20-positive mitochondrial objects, n=6. (D) Representative images of efferocytosis by IL-4-treated WT and lipin-1mKO BMDMs following the indicated siRNA treatments. CFSE-labeled apoptotic cells are shown in green. Scale bar, 50 μm. (E) Quantification of efferocytosis, expressed as the percentage of macrophages containing at least one apoptotic cell, n=4. Significance was assessed by one-way ANOVA. \**p* < 0.05, \*\**p* < 0.01, \*\*\**p* < 0.001, \*\*\*\**p*< 0.0001; ns, not significant.

Together, these findings indicate that experimentally inducing mitochondrial elongation by DRP1 depletion in WT BMDMs does not reproduce the lipin-1-dependent defects in efferocytosis under the conditions tested, and that inducing fragmentation by OPA1 depletion in lipin-1mKO BMDMs does not rescue it. Thus, altered mitochondrial morphology is associated with lipin-1 deficiency but is unlikely to be the principal cause of its efferocytic phenotype. These findings prompted us to examine alternative mechanisms by which lipin-1 regulates efferocytosis and mitochondrial function, including homeostasis of mitochondrial membrane phospholipids.

### Lipin-1 deficiency preferentially reduces mitochondrial inner membrane proteins and alters cristae architecture in IL-4-stimulated macrophages

Because altering mitochondrial morphology did not reproduce or rescue the lipin-1-dependent defects in efferocytosis, we next asked whether lipin-1 deficiency was associated with broader changes in mitochondrial protein composition. We isolated mitochondria-enriched fractions from IL-4-treated WT and lipin-1mKO BMDMs and analyzed protein abundance by LC–MS/MS. Immunoblotting showed TOM-20 enrichment relative to the cytosolic protein GAPDH in the mitochondrial fraction, confirming mitochondrial enrichment before proteomic analysis (Figure 4A).

**Figure 4.**
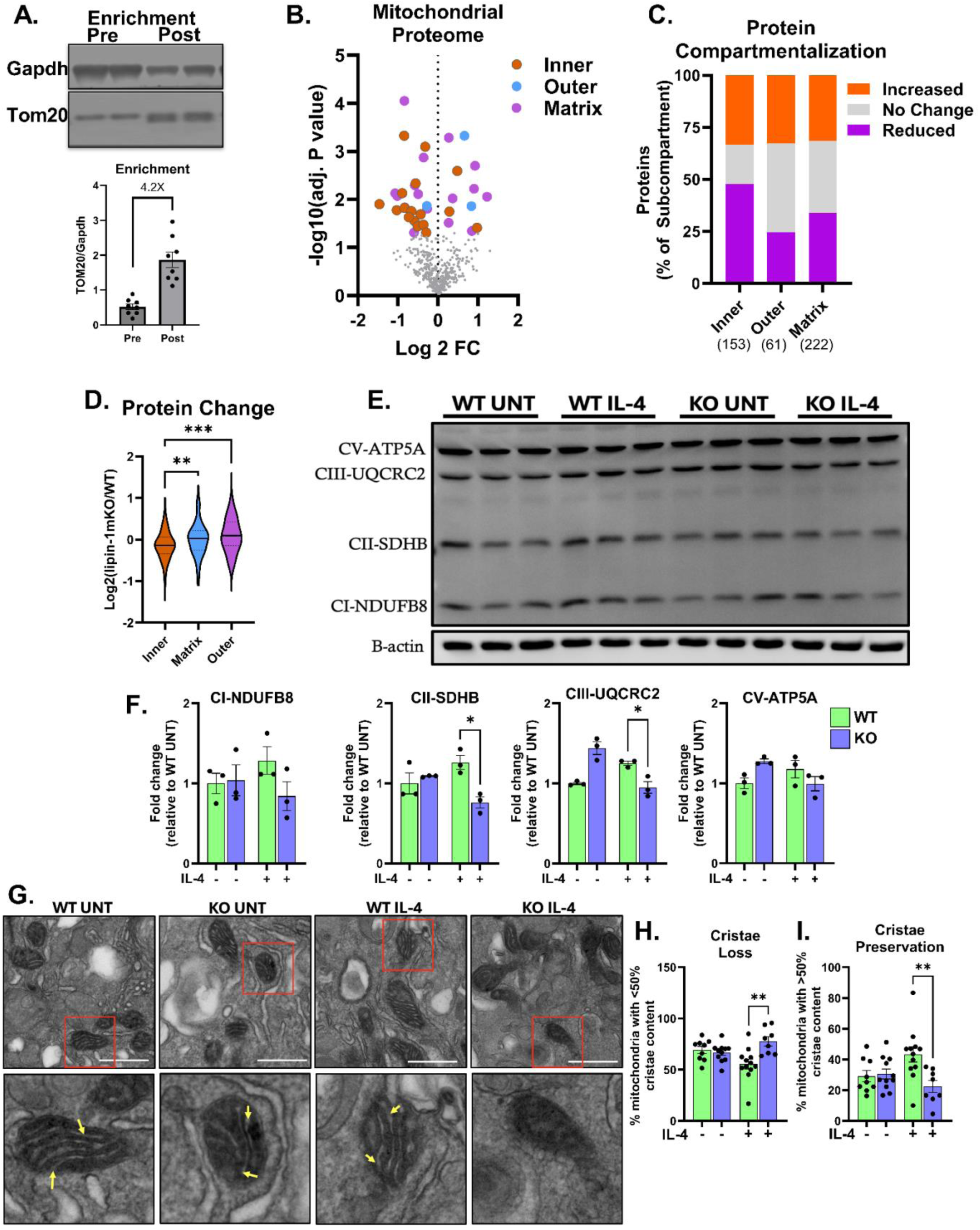
Lipin-1 deficiency preferentially reduces mitochondrial inner-membrane proteins in IL-4-stimulated macrophages. Mitochondria-enriched fractions from IL-4-treated WT and lipin-1mKO BMDMs were analyzed by LC-MS/MS. (A) Representative immunoblot demonstrating enrichment of the mitochondrial protein TOM-20 and depletion of the cytosolic protein GAPDH in the mitochondria-enriched fraction relative to whole-cell lysate. (B) Volcano plot of mitochondrial protein abundance in IL-4-treated lipin-1mKO relative to WT BMDMs. Proteins were assigned to mitochondrial compartments using MitoCarta 3.0 annotations. A total of 436 proteins were quantified, including 222 matrix, 153 mitochondrial inner-membrane (MIM), and 61 mitochondrial outer-membrane (MOM) proteins. Thirty-six proteins differed significantly between genotypes by two-sided unpaired *t*-test (*P* < 0.05), including 24 decreased and 12 increased proteins. (C) Distribution of MIM, matrix, and MOM proteins classified as decreased (>10% reduction), unchanged (within ±10%), or increased (>10% increase) in lipin-1mKO relative to WT BMDMs. The distribution differed among mitochondrial compartments (Pearson χ²(4) = 18.88, P = 8.31 × 10⁻⁴). Pairwise comparisons of the proportion of decreased versus not-decreased proteins were performed using two-sided Fisher’s exact tests with Holm correction. (D) Distribution of log2 fold changes for proteins assigned to the matrix, MIM, or MOM. Distributions were compared using a Kruskal-Wallis test followed by pairwise two-sided Mann-Whitney U tests with Holm correction. (E) Representative immunoblot of OXPHOS complex I (NDUFB8), complex II (SDHB), complex III (UQCRC2), and complex V (ATP5A) subunits in untreated and IL-4-treated WT and lipin-1mKO BMDMs. (F) Densitometric quantification of the indicated OXPHOS proteins normalized to the loading control (β-Actin), and fold change was determined relative to the untreated WT group. (G) Representative electron micrographs showing cristae morphology. Scale bar, 600nm. Bottom panels are zoomed in pictures of the regions in the red boxes. Yellow arrows indicate intact cristae. (H) Quantification of mean percentage of mitochondria with less than 50% visible intact cristae. (I) Quantification of mean percentage of mitochondria with more than 50% visible intact cristae. Each point represents an independent BMDM preparation, n = 3 per group. Data are mean ± SEM. Statistical analysis for (F): one-way ANOVA. \**p* < 0.05, \*\**p* < 0.01, \*\*\**p* < 0.001, \*\*\*\**p*< 0.0001; ns, not significant.

MitoCarta 3.0 annotations were used to identify mitochondrial proteins and assign their predominant localization to the matrix, mitochondrial inner membrane (MIM), or mitochondrial outer membrane (MOM). We quantified 436 proteins, including 222 matrix, 153 MIM, and 61 MOM proteins, of which 36 were significantly different: 24 decreased and 12 increased (Figure 4B). Among the decreased, 14 of these proteins localized to the MIM. In comparison, 16 matrix proteins met this threshold, with 9 decreased and 7 increased, whereas only 3 MOM proteins were identified, with 1 decreased and 2 increased. The reduced MIM proteins represented several mitochondrial functions rather than a single protein complex. Six complex I subunits were decreased (NDUFA11, NDUFS8, NDUFS5, NDUFA5, NDUFC2, and NDUFB7), together with the complex II subunit SDHC and the complex III subunit UQCR10. Other reduced MIM proteins included COQ8B, which participates in CoQ metabolism; RMND1, which supports mitochondrial translation and OXPHOS biogenesis; MICOS13, a component of the cristae-organizing MICOS complex; CPOX and SFXN2, which are associated with mitochondrial heme and iron metabolism; and the mitochondrial Na^+^/Ca^2+^ exchanger SLC8B1. Thus, the MIM proteins reduced in lipin-1mKO BMDMs encompassed respiratory-chain components as well as proteins involved in mitochondrial membrane organization and metabolism.

The predominance of decreased MIM proteins among the proteins identified prompted us to ask whether this pattern extended across the mitochondrial proteome rather than being limited to proteins meeting the p-value threshold. Because mitochondrial proteins, particularly respiratory-chain components, function within stoichiometrically coordinated multiprotein complexes, modest but coordinated changes in protein abundance may be biologically informative even when individual proteins do not exhibit large fold changes ^32^. We therefore used a ±10% threshold as an operational cutoff to classify proteins as decreased, unchanged, or increased and determine whether directional changes were preferentially associated with specific mitochondrial subcompartments. The distribution of these categories differed significantly among mitochondrial compartments (Pearson χ²(4) = 18.88, P = 8.31 × 10⁻⁴; (Figure 4C). MIM proteins showed a significant change in frequency of decreased abundance, with 73 of 153 proteins (47.7%) classified as decreased, compared with 75 of 222 matrix proteins (33.8%) and 15 of 61 MOM proteins (24.6%). In contrast, approximately one-third of the proteins in each compartment were unchanged. Consistent with this categorical analysis, visualization of log2 fold changes for all quantified proteins showed a broader downward shift in the MIM protein distribution relative to matrix and MOM proteins (Figure 4D). Together, these analyses demonstrate that reduced protein abundance is preferentially associated with the mitochondrial inner membrane rather than reflecting a uniform loss of mitochondrial proteins.

To determine whether this broader MIM phenotype was also reflected in respiratory-chain protein abundance, we quantified representative subunits of OXPHOS complexes I, II, III, and V by immunoblotting (Figure 4E, F). Following IL-4 stimulation, lipin-1mKO BMDMs exhibited significantly reduced abundance of SDHB and UQCRC2, representing complexes II and III, respectively, whereas NDUFB8 and ATP5A did not differ significantly between genotypes. These findings provide independent support for altered respiratory-chain protein abundance in lipin-1-deficient macrophages. Together, the proteomic and immunoblotting data indicate that lipin-1 deficiency is associated with a preferential reduction in mitochondrial inner-membrane proteins, including components of the respiratory machinery.

Given that cristae membranes provide the structural framework that organizes and concentrates MIM proteins, including OXPHOS complexes, we next investigated whether the reduction in abundance of MIM protein was accompanied by alterations in cristae architecture. Cristae morphology influences the spatial organization, assembly, stability of MIM proteins and complexes, and contributes to efficient protein import and oxidative phosphorylation ^33^. In response to IL-4 stimulation, lipin-1mKO BMDMs had significantly altered cristae integrity, characterized by a higher percentage of mitochondria having fewer than 50% visible intact cristae compared with WT BMDMs (Figure 4G-4I). These findings suggest that lipin-1 deficiency disrupts not only the MIM proteome but also the ultrastructural organization required to maintain it. This broad inner-membrane phenotype and altered cristae architecture prompted us to examine whether lipin-1 deficiency also alters the mitochondrial phospholipid environment that supports inner-membrane organization and cristae architecture

### Lipin-1 deficiency reduces cardiolipin abundance in IL-4-stimulated macrophages

Cardiolipin is critical for the formation and maintenance of cristae, the stabilization and functional organization of OXPHOS complexes and assembly of their higher-order super-complexes, thereby optimizing electron transport and ATP synthesis ^34–36^ (Figure 5A). Because lipin-1 deficiency was associated with reduced abundance of OXPHOS subunits and mitochondrial inner-membrane proteins, and altered cristae morphology, we investigated whether lipin-1 also regulates cardiolipin abundance. Cardiac-specific lipin-1 deficiency was previously shown to increase PA abundance while paradoxically reducing cardiolipin and mitochondrial respiration in mouse heart tissues, suggesting that cardiolipin homeostasis depends on more than total cellular PA availability ^20^. We initially assessed cardiolipin-associated fluorescence in untreated and IL-4-treated WT and lipin-1mKO BMDMs using NAO staining followed by widefield fluorescence microscopy and flow cytometry. Both approaches showed reduced NAO fluorescence in IL-4-treated lipin-1mKO BMDMs compared with WT BMDMs (Figure 5B–E). Because NAO fluorescence can be influenced by mitochondrial membrane potential, respiratory state, and membrane organization, we directly quantified cardiolipin by LC–MS/MS. This analysis confirmed a significant reduction in total cardiolipin abundance in IL-4-treated lipin-1mKO BMDMs (Figure 5F). These findings are consistent with previous observations that cardiac-specific lipin-1 deficiency increases PA abundance while reducing cardiolipin content and mitochondrial respiration. Given that PA is both a substrate of lipin-1 PAP activity and a precursor for cardiolipin biosynthesis, the reduction in cardiolipin following lipin-1 deficiency raises questions regarding the intracellular distribution and metabolic fate of PA. Lipin-1 may therefore influence cardiolipin homeostasis through mechanisms involving PA compartmentalization, mitochondrial delivery, or utilization, as well as cardiolipin remodeling or turnover.

**Figure 5.**
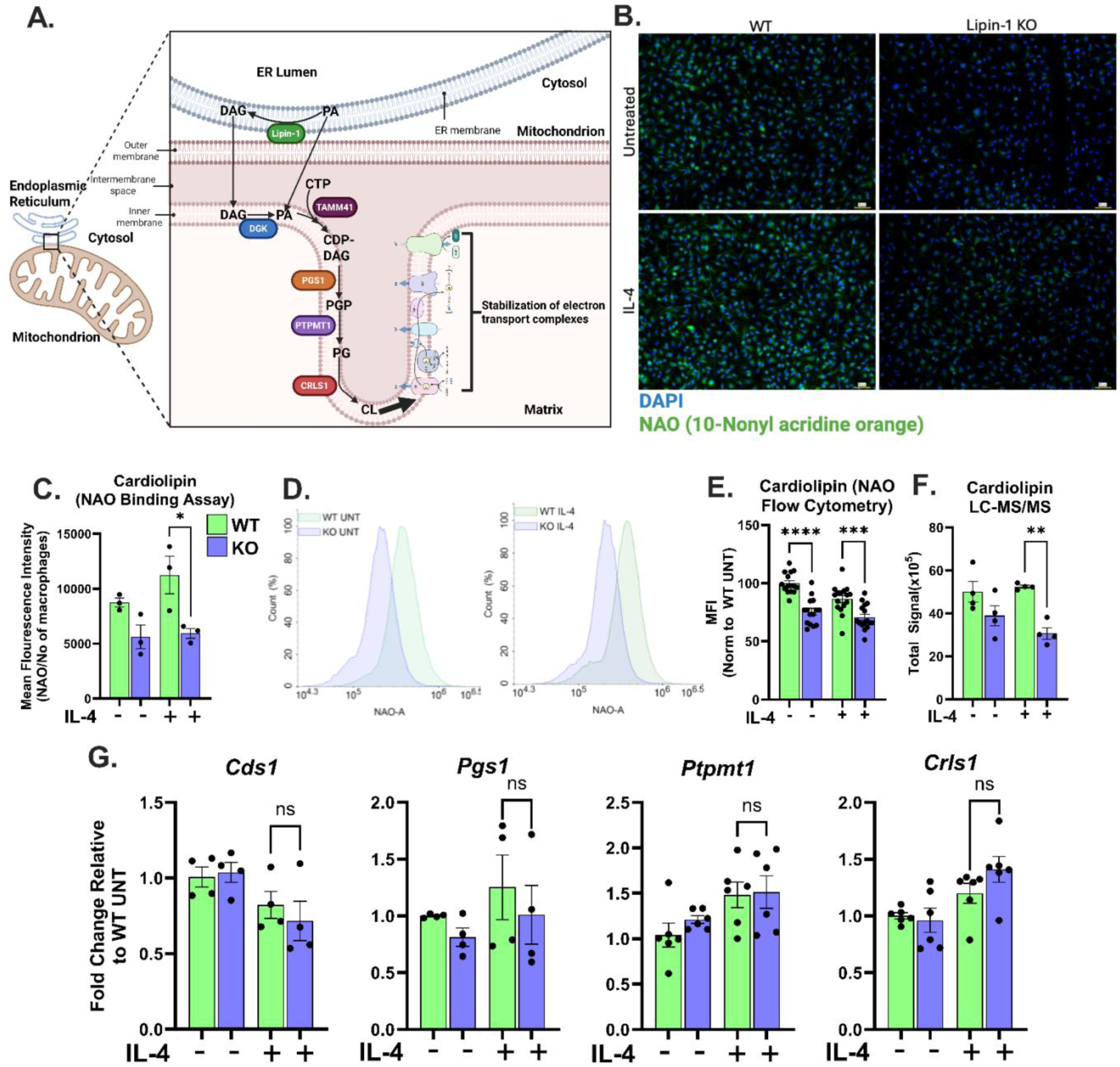
Lipin-1 deficiency reduces cardiolipin abundance in IL-4-stimulated macrophages. (A) Schematic illustrating mitochondrial cardiolipin biosynthesis and the relationship between phosphatidic acid and lipin-1 to mitochondrial phospholipid metabolism. (B, C) Cardiolipin-associated fluorescence was assessed using nonyl acridine orange (NAO) in untreated or IL-4-treated (40 ng/mL, 6 h) WT and lipin-1mKO BMDMs. (B) Representative fluorescence microscopy images of NAO staining (green); nuclei are shown in blue. Scale bar, 50 μm. (C) Quantification of NAO fluorescence by microscopy, expressed as integrated fluorescence intensity normalized to cell number. (D) Representative flow-cytometry histograms of NAO fluorescence in untreated and IL-4-treated WT and lipin-1mKO BMDMs. (E) Quantification of NAO median fluorescence intensity (MFI), normalized to untreated WT BMDMs. (F) Relative total cardiolipin abundance measured directly by targeted LC-MS/MS analysis and normalized to the untreated WT group n=4. (G) qRT-PCR analysis of *Cds1, Pgs1, Ptpmt1,* and *Crls1* expression in untreated and IL-4-treated WT and lipin-1mKO BMDMs. Transcript abundance was normalized to *Gapdh* and expressed relative to untreated WT BMDMs. Each point represents an independent BMDM preparation, n = 4-6 per group. Data are shown as means ± SEM. Statistical analyses were done using one-way ANOVA. \**p* < 0.05, \*\**p* < 0.01, \*\*\**p* < 0.001, \*\*\*\**p*< 0.0001; ns, not significant.

Because lipin-1 PAP activity consumes PA, the reduction in cardiolipin is not readily explained by decreased total PA availability. Instead, the findings suggest that lipin-1 may regulate cardiolipin abundance through an unresolved mechanism involving PA compartmentalization, mitochondrial delivery, utilization, or cardiolipin stability.

We next determined whether reduced cardiolipin abundance was associated with altered expression of genes involved in cardiolipin synthesis, again to confirm if lipin-1 transcriptional co-regulatory activity could be involved. The transcript abundance of *Cds1*, *Pgs1*, *Ptpmt1*, and *Crls1* did not differ significantly between IL-4-treated WT and lipin-1mKO BMDMs (Figure 5G). These findings indicate that reduced cardiolipin abundance in lipin-1mKO macrophages is not explained by altered steady-state expression of the selected cardiolipin synthesis and remodeling genes. Together, these findings demonstrate that lipin-1 deficiency reduces cardiolipin abundance in IL-4-stimulated macrophages without broadly altering the expression of selected cardiolipin-pathway genes. Thus, lipin-1 appears to regulate cardiolipin homeostasis through a mechanism beyond transcriptional control of these enzymes. These observations prompted us to determine whether cardiolipin is functionally required for lipin-1-dependent mitochondrial respiration and efferocytosis.

### Cardiolipin homeostasis supports IL-4-induced mitochondrial respiration and efferocytosis

Because lipin-1 deficiency reduced cardiolipin abundance, we next tested whether disrupting cardiolipin synthesis was sufficient to impair IL-4-induced macrophage respiration and efferocytosis. WT BMDMs were transfected with either scrambled or *Ptpmt1*-targeting siRNA (si-Ptpmt1) to reduce PTPMT1. PTPMT1 catalyzes the conversion of phosphatidylglycerophosphate to phosphatidylglycerol, the penultimate step in de novo cardiolipin synthesis. Immunoblotting confirmed reduction in PTPMT1 protein abundance (Figure 6A). We then treated these cells with 40 ng/mL IL-4, after which we conducted the efferocytosis assay on the BMDMs. Compared with WT treated with scrambled siRNA, the si-Ptpmt1-transfected WT BMDMs reduced efferocytosis to a level not significantly different from control-transfected lipin-1mKO BMDMs. (Figure 6B). These findings are consistent with a requirement for intact cardiolipin biosynthesis in macrophage efferocytosis.

**Figure 6.**
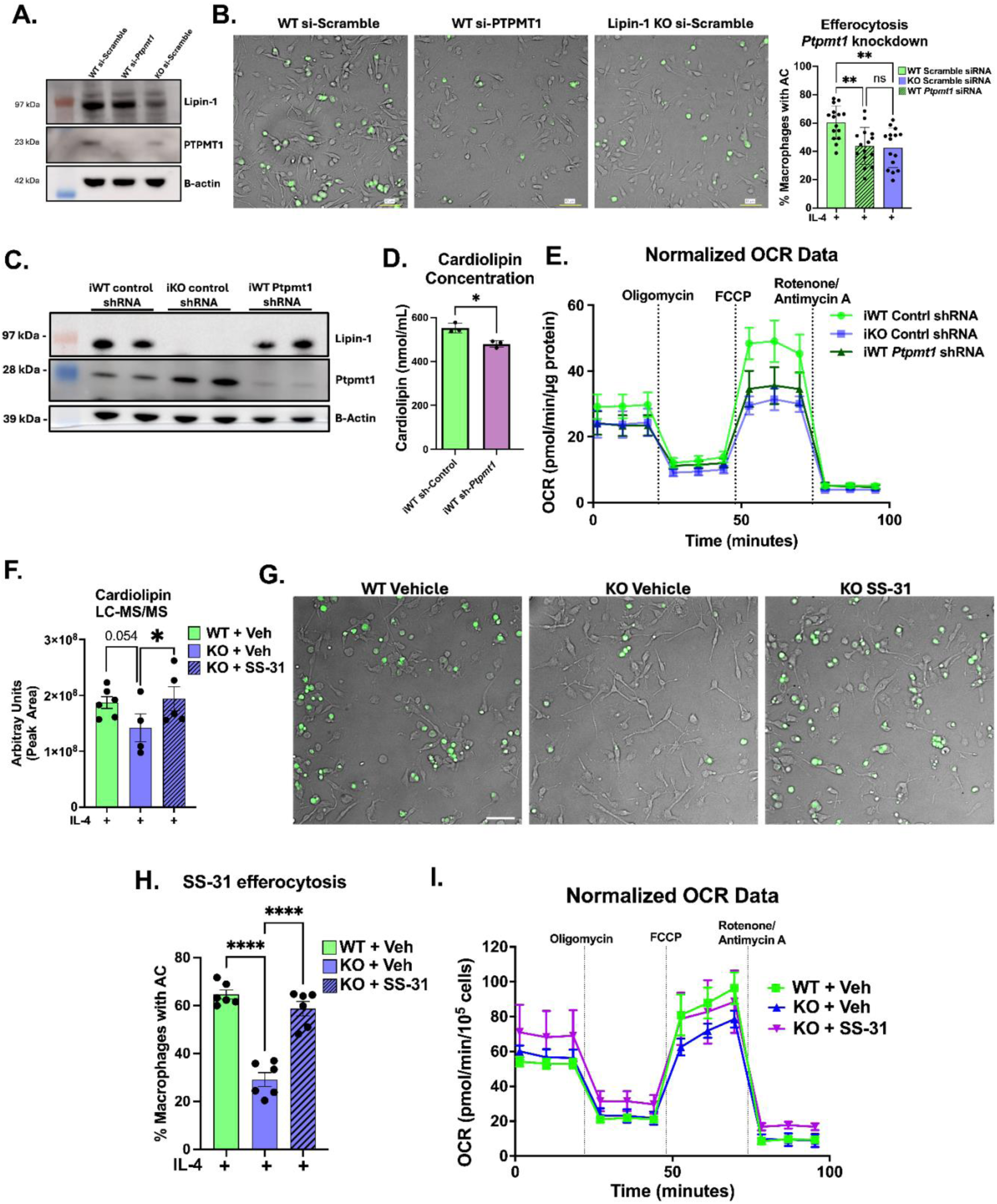
Cardiolipin homeostasis supports IL-4-induced mitochondrial respiration and macrophage efferocytosis. (A, B) WT BMDMs were transfected with control or *Ptpmt1*-targeting siRNA; control-transfected lipin-1mKO BMDMs were included for comparison. (A) Representative immunoblot confirming depletion of PTPMT1. β-Actin served as the loading control. (B) Representative images and quantification of efferocytosis following IL-4 stimulation. CFSE-labeled apoptotic cells are shown in green. Efferocytosis was calculated as the percentage of macrophages containing at least one apoptotic cell. Scale bar, 50 μm. (C, D) Immortalized WT BMDMs (iWT) were transduced with lentiviral vectors expressing doxycycline-inducible control or *Ptpmt1* shRNA, while immortalized lipin-1 KO BMDMs (iKO) were transduced with control shRNA vector. (C) Representative immunoblot confirming depletion of PTPMT1. (D) Cardiolipin fluorometric assay of iWT expressing doxycycline-inducible control or *Ptpmt1* shRNA. (E) Mitochondrial respiration was measured by Seahorse XF Mito Stress Test following IL-4 stimulation. OCR was measured during sequential addition of oligomycin, FCCP, and rotenone plus antimycin A. (F-H) Lipin-1mKO BMDMs were pretreated with SS-31 (0.5 μM) for 2 h and subsequently stimulated with 40 ng/mL IL-4 for 6 h in the continued presence of SS-31; WT and lipin-1mKO vehicle-treated BMDMs served as controls. (F) Relative cardiolipin abundance measured by LC-MS/MS. (G, H) Representative images and quantification of efferocytosis. Scale bar, 50 μm. Each point represents an independent BMDM preparation, n = 6 per group. (I) Lipin-1mKO BMDMs were pretreated with SS-31 (10 μM) for 2 h and subsequently stimulated with IL-4 in the continued presence of SS-31. OCR was measured by the Seahorse XF Mito Stress Test. Bar graphs are shown as means ± SEM. Statistical analyses were done using unpaired *t* test (D) one-way ANOVA. \**p* < 0.05, \*\**p* < 0.01, \*\*\**p* < 0.001, \*\*\*\**p*< 0.0001; ns, not significant.

Given the importance of cardiolipin in maintaining respiratory chain function, we next examined whether disruption of cardiolipin biosynthesis recapitulated the reduction in OXPHOS observed in lipin-1mKO macrophages. To test this using an independent genetic approach, we generated immortalized WT and lipin-1mKO bone marrow-derived macrophages (iBMDMs) using the established Cre-J2 retroviral immortalization method. J2-immortalized macrophages have been shown to retain many functional and metabolic properties of primary BMDMs ^37^. We first confirmed that WT and lipin-1mKO iBMDMs recapitulated the phenotype-dependent differences in IL-4-induced efferocytosis and OXPHOS observed in primary BMDMs (Figure S2). This system also enabled stable introduction of doxycycline-inducible constructs to deplete PTPMT1, complementing transient siRNA treatment in primary BMDMs and avoiding additional transfection or nucleofection immediately before metabolic analysis. Doxycycline-induced depletion of PTPMT1 in WT iBMDMs was confirmed by immunoblotting (Figure 6C) while cardiolipin levels were measured by cardiolipin fluorometric assay (Figure 6D). We then assessed mitochondrial respiration using the Seahorse XF Cell Mito Stress Test. PTPMT1 depletion in WT iBMDMs resulted in an impaired OXPHOS (Figure 6E). Collectively, these findings demonstrate that disrupting cardiolipin biosynthesis recapitulates both the efferocytosis and mitochondrial respiration defects observed in lipin-1-deficient macrophages, supporting cardiolipin homeostasis as a component of lipin-1-dependent regulation of mitochondrial biology and macrophage pro-resolving function.

We next tested whether pharmacologically targeting cardiolipin-associated mitochondrial dysfunction could rescue efferocytosis in lipin-1mKO BMDMs. SS-31 (elamipretide) is a mitochondria-targeted peptide that interacts with cardiolipin, stabilizing it from oxidative damage and supporting cardiolipin-dependent cristae and respiratory function ^38,39^. Lipin-1mKO BMDMs were pretreated with SS-31 for 2 hours and subsequently stimulated with IL-4 in the continued presence of SS-31 for 6 hours. LC-MS/MS demonstrated that SS-31 increased cardiolipin abundance in lipin-1mKO BMDMs relative to vehicle-treated lipin-1mKO controls, restoring cardiolipin to a level not significantly different from vehicle-treated WT BMDMs (Figure 6F). SS-31 treatment significantly increased efferocytosis in lipin-1mKO BMDMs compared with vehicle-treated lipin-1mKO BMDMs, restoring efferocytic activity to a level not significantly different from vehicle-treated WT BMDMs (Figure 6G & 6H). SS-31 treatment also increased basal and maximal respiration in lipin-1mKO BMDMs compared with vehicle-treated lipin-1mKO BMDMs. These respiratory parameters did not differ significantly from those measured in vehicle-treated WT BMDMs (Figure 6I). Together, PTPMT1 depletion and SS-31 treatment provide complementary evidence that cardiolipin homeostasis supports efferocytosis and mitochondrial respiration in IL-4-treated macrophages. These findings further suggest that reduced cardiolipin contributes to but may not fully account for the bioenergetic and efferocytic defects caused by lipin-1 deficiency.

### SS-31 reduces apoptotic thymocyte accumulation in lipin-1mKO mice following dexamethasone-induced thymic injury

We previously demonstrated that myeloid lipin-1 supports macrophage efferocytosis and resolution-associated outcomes in multiple in vivo models ^8,9,40^. Because SS-31 restored cardiolipin abundance, mitochondrial respiration, and efferocytosis in lipin-1mKO BMDMs in vitro, we next tested whether SS-31 improved apoptotic-cell clearance in vivo. We used the dexamethasone-induced thymic involution model, in which extensive thymocyte apoptosis creates a high apoptotic-cell burden that thymic macrophages must continually clear. Lipin-1mKO mice received a single intraperitoneal injection of SS-31 (5 mg/kg) or vehicle, whereas littermate WT controls received vehicle. Six hours later, mice were injected intraperitoneally with 250 µg dexamethasone in 250 µL PBS, and thymi were collected 18 hours after dexamethasone administration (Figure 7A) ^41–43^. One thymic lobe was processed for flow cytometry and total-cell enumeration, and the contralateral lobe was fixed and paraffin-embedded for histologic analysis. Vehicle-treated lipin-1mKO mice had more ApoTracker-positive thymic cells and greater total thymic cellularity than vehicle-treated WT mice after dexamethasone administration (Figure 7B, C). SS-31 treatment reduced both apoptotic-cell burden and thymic cellularity in lipin-1mKO mice. These endpoints did not differ significantly from those measured in vehicle-treated WT mice. To determine whether the reduced apoptotic-cell burden reflected enhanced macrophage efferocytosis, thymic sections were stained with TUNEL to identify apoptotic cells and Mac2 to identify macrophages. In situ efferocytosis was quantified by distinguishing TUNEL-positive cells associated with Mac2-positive macrophages from free TUNEL-positive cells and calculating the macrophage-associated: free apoptotic-cell ratio (Figure 7D, E). Together, these findings demonstrate that SS-31 reduces apoptotic cell accumulation and increases the macrophage-associated:free apoptotic cell ratio in lipin-1mKO thymi following dexamethasone-induced injury. These results indicate that SS-31 improves apoptotic thymocyte clearance in lipin-1mKO mice and support the hypothesis that cardiolipin-associated mitochondrial dysfunction contributes to defective efferocytosis caused by myeloid lipin-1 deficiency. Because SS-31 was administered systemically, the experiment does not by itself establish that the rescue is exclusively macrophage-intrinsic.

**Figure 7.**
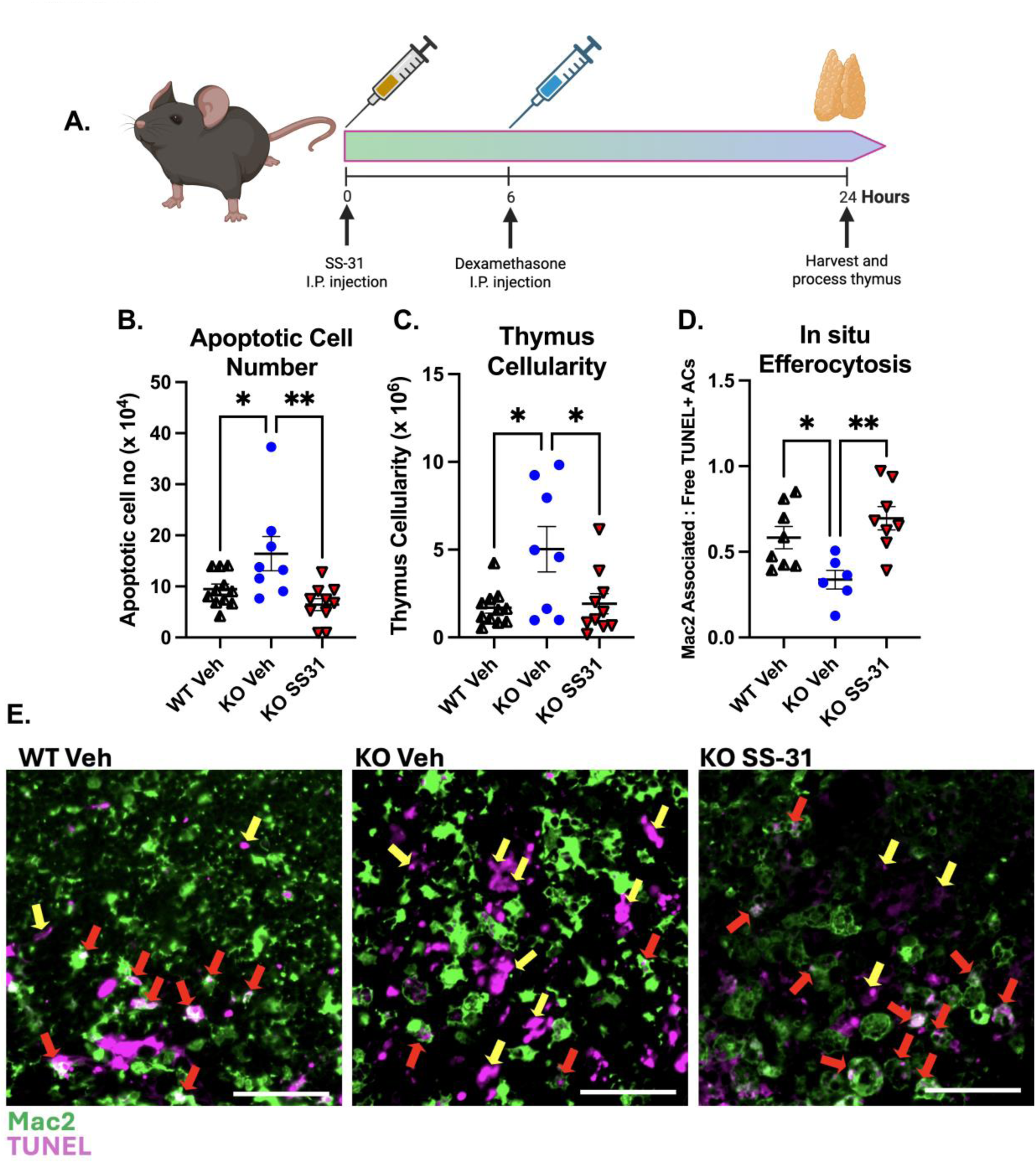
SS-31 improves apoptotic-cell clearance in lipin-1mKO mice following dexamethasone-induced thymic injury. (A) Experimental design. WT mice received vehicle, whereas lipin-1mKO mice received vehicle or SS-31 (5 mg/kg, intraperitoneally). Six hours later, mice received dexamethasone (250 μg, intraperitoneally), and thymi were collected 18 h after dexamethasone administration. (B) Number of ApoTracker-positive apoptotic thymic cells determined by flow cytometry. (C) Total thymic cellularity. (D) Quantification of in situ efferocytosis; the ratio of macrophage-associated to free TUNEL-positive cells. (E) Representative thymic sections stained for TUNEL-positive apoptotic cells (ACs) and Mac-2-positive macrophages; red arrows indicate macrophage-associated ACs and yellow arrows indicate free TUNEL-positive ACs. Scale bar, 50 μm. Each point represents one mouse. Data are presented as mean ± SEM. Statistical analyses were done using one-way ANOVA. \**p* < 0.05, \*\**p* < 0.01, \*\*\**p* < 0.001, \*\*\*\**p*< 0.0001; ns, not significant.

## DISCUSSION

Macrophage efferocytosis and inflammation resolution depend on coordinated metabolic adaptation, with a greater reliance on mitochondrial oxidative metabolism ^3,7,8^. In our previous studies, we found that macrophage lipin-1 supports OXPHOS, efferocytosis, inflammation resolution, and atheroprotection, independent of lipin-1 PAP activity ^7–9^. However, how lipin-1 supports mitochondrial function remained unclear. In the present study, lipin-1-deficient macrophages showed reduced cardiolipin abundance, altered mitochondrial morphology, and lower levels of mitochondrial inner-membrane proteins and electron transport chain components. We also found that disrupting cardiolipin synthesis impaired macrophage efferocytosis, whereas treatment of lipin-1-deficient macrophages with the cardiolipin-interacting peptide SS-31 improved OXPHOS and efferocytosis. These findings support a functional role for cardiolipin homeostasis in the lipin-1-dependent mitochondrial program that promotes macrophage efferocytosis, while the mechanism by which lipin-1 maintains cardiolipin abundance remains unresolved.

One potential explanation for the metabolic effects of lipin-1 was its established transcriptional co-regulatory activity. Nuclear lipin-1 interacts with transcriptional regulators including Peroxisome Proliferator-Activated Receptor Gamma Coactivator 1α (PGC-1α) to promote oxidative metabolic programs and can repress SREBP-dependent lipogenic transcription ^21–23^. We therefore considered whether the impaired β-oxidation, altered lipid handling, and reduced mitochondrial respiration previously observed in lipin-1-deficient macrophages could result from broad changes in metabolic gene expression. However, bulk RNA sequencing and targeted gene expression analyses did not reveal widespread suppression of canonical PPAR-or SREBP-associated programs or substantial differences in the expression of the β-oxidation and ceramide-synthesis genes examined. These findings argue against broad transcriptional regulation of these metabolic pathways as the principal explanation for the lipin-1-dependent respiratory and efferocytic phenotypes. They do not, however, exclude more selective transcriptional effects of lipin-1. Indeed, enrichment of IRF1-and IRF8-associated gene programs suggests that lipin-1 deficiency may influence more focused immunometabolic transcriptional pathways, consistent with emerging roles for these transcription factors in mitochondrial function and cellular metabolism ^44–47^.

Lipin-1 deficiency was also associated with a more elongated mitochondrial network, raising the possibility that altered mitochondrial morphology contributes to the impaired metabolic and efferocytic phenotype. Mitochondrial fission has previously been linked to macrophage efferocytosis, particularly during successive rounds of apoptotic-cell uptake ^26^. However, reciprocal manipulation of mitochondrial morphology did not reproduce or rescue the lipin-1-dependent efferocytosis phenotype under the conditions examined here. OPA1 depletion, which promotes mitochondrial fragmentation, was insufficient to restore efferocytosis in lipin-1-deficient macrophages, whereas DRP1 depletion, which promotes elongation, did not reproduce the defect in WT cells. Thus, although lipin-1 deficiency is associated with altered mitochondrial morphology, mitochondrial elongation alone appears insufficient to account for the loss of macrophage efferocytic capacity. Because our measurements relied primarily on static mitochondrial morphology and because DRP1 and OPA1 regulate processes beyond fission and fusion, including mitophagy, cristae organization, respiratory complex function, apoptosis, and mitochondrial–ER communication, these experiments do not exclude contributions of mitochondrial dynamics or quality-control pathways to lipin-1-dependent macrophage function ^48–55^.

The inability of mitochondrial fission-fusion dynamics alone to account for the efferocytosis defect suggests that other aspects of mitochondrial biology may contribute to lipin-1-dependent macrophage function. Notably, our proteomic analysis of enriched mitochondrial fractions revealed a preferential reduction in proteins localized to the mitochondrial inner membrane (MIM) in lipin-1mKO BMDMs stimulated by IL-4. Nearly half of MIM proteins were reduced by more than 10% in lipin-1mKO BMDMs (47.7%), compared with 33.8% of matrix proteins and 24.6% of outer-membrane proteins, and MIM proteins were significantly more likely to be decreased than proteins in either of the other compartments. The proteins contributing to this pattern spanned multiple inner-membrane functions. Reduced proteins included several complex I subunits, as well as components of complexes II and III, OXPHOS assembly and mitochondrial translation machinery, the cristae-associated protein MICOS13, and proteins involved in metabolite transport and mitochondrial heme/iron metabolism. Thus, the phenotype was not restricted to a single respiratory complex or pathway. Consistent with the proteomic findings, immunoblotting showed reduced abundance of the complex II subunit SDHB and the complex III subunit UQCRC2 in IL-4-treated lipin-1-deficient macrophages, although not all representative OXPHOS subunits were significantly reduced. These findings extend our previous observations that lipin-1-deficient macrophages have impaired mitochondrial respiration and altered TCA-cycle metabolism ^7^ and are consistent with the reduced abundance of SDHA previously observed in lipin-1-deficient peritoneal macrophages during zymosan-induced inflammation resolution ^8^. Because respiratory-chain integrity influences not only ATP production but also metabolites and signaling pathways that regulate macrophage phenotype, including succinate metabolism, HIF-1α stabilization, and IL-10 production, this broader inner-membrane defect could have consequences beyond bioenergetics alone ^56–58^. Importantly, the compartment-specific nature of the proteomic shift is difficult to explain by a generalized reduction in mitochondrial content and instead points to disruption of a property of the inner mitochondrial membrane that supports the stability or maintenance of multiple classes of proteins. In line with this interpretation, IL-4-treated lipin-1-deficient macrophages had significantly altered cristae assembly and structural integrity, providing addition evidence of mitochondrial inner membrane abnormalities that may contribute to the reduced abundance of MIM proteins ^33^.

Cardiolipin represents one candidate determinant of this inner membrane and cristae phenotype. Cardiolipin is highly enriched in the mitochondrial inner membrane and contributes to cristae organization, respiratory-complex and supercomplex stability, mitochondrial protein import, and carrier-protein function ^11,35,36,59^. This is particularly relevant to our proteomic findings, because the proteins reduced in lipin-1-deficient macrophages were not confined to a single respiratory complex but included multiple respiratory-chain components, metabolite transporters, and the cristae-associated protein MICOS13. We found that cardiolipin abundance was reduced in IL-4-stimulated lipin-1-deficient macrophages, extending previous observations that cardiac lipin-1 deficiency is associated with reduced cardiolipin and impaired mitochondrial respiration ^20^. The relationship between lipin-1 and cardiolipin is particularly intriguing given the established enzymatic function of lipin-1 in phospholipid metabolism. Lipin-1 converts phosphatidic acid (PA), an upstream precursor of cardiolipin biosynthesis, to diacylglycerol (DAG). Based on this enzymatic activity, loss of lipin-1 would be expected to favor PA accumulation, potentially increasing substrate availability for cardiolipin biosynthesis rather than reducing cardiolipin abundance. Moreover, lipin-1 deficiency did not reduce the expression of the cardiolipin biosynthetic genes examined, and the mitochondrial proteome did not reveal a coordinated loss of cardiolipin biosynthetic proteins ^60–62^. Together, these observations suggest that the reduction in cardiolipin cannot be readily explained by the established phosphatidic acid phosphatase activity of lipin-1 or by a broad reduction in cardiolipin biosynthetic machinery. Instead, lipin-1 may influence cardiolipin homeostasis through mechanisms involving intracellular lipid compartmentalization, precursor trafficking or utilization, or cardiolipin remodeling and turnover.

How lipin-1 maintains macrophage cardiolipin abundance therefore remains an important unresolved question. Total cellular phosphatidic acid abundance may not reflect the pool of PA available to mitochondria, and cardiolipin abundance can be influenced by substrate compartmentalization, phospholipid trafficking, biosynthesis, remodeling, oxidation, and turnover. Potential mechanisms could therefore include altered phospholipid exchange at mitochondrial-ER contact sites, changes in PA delivery to mitochondria, regulation of lipid-transfer pathways such as TRIAP1-PRELID, or effects on cardiolipin remodeling or stability ^63–66^. The current study does not distinguish among these possibilities. Instead, it establishes the previously unrecognized association between macrophage lipin-1 and cardiolipin abundance and provides a rationale for determining how lipin-1 influences mitochondrial phospholipid homeostasis in future studies.

Importantly, our data also provide functional support for a relationship between cardiolipin homeostasis and macrophage efferocytosis. Depletion of PTPMT1, which participates in de novo cardiolipin synthesis, reduced cardiolipin abundance and impaired efferocytosis in WT macrophages. Conversely, treatment of lipin-1-deficient macrophages with SS-31 increased cardiolipin abundance and improved mitochondrial respiration and efferocytosis. SS-31 interacts with cardiolipin and has been reported to stabilize cardiolipin-associated mitochondrial membrane organization and limit cardiolipin oxidation ^39,67^. The complementary effects of disrupting cardiolipin synthesis and supporting cardiolipin-associated mitochondrial function therefore suggest that reduced cardiolipin contributes to the metabolic and efferocytic phenotype of lipin-1-deficient macrophages. Nevertheless, neither PTPMT1 depletion nor SS-31 treatment is uniquely specific to cardiolipin, and these experiments do not demonstrate that cardiolipin loss is the sole cause of the lipin-1-deficient phenotype. Rather, they support cardiolipin homeostasis as one important component of the mitochondrial program required for efficient macrophage respiration and apoptotic-cell clearance.

The potential relevance of this pathway in vivo is supported by our findings in dexamethasone-induced thymic injury. Myeloid-specific lipin-1 deficiency has previously been associated with impaired inflammation resolution across multiple models, including delayed neutrophil clearance during zymosan-induced peritonitis, impaired wound healing, and increased atherosclerotic plaque burden and necrotic-core formation ^8,9,40^. Following dexamethasone-induced thymic injury, lipin-1-deficient mice similarly exhibited increased accumulation of apoptotic thymocytes, whereas SS-31 treatment reduced apoptotic-cell burden and thymic cellularity. These observations are consistent with improved apoptotic-cell clearance and support the potential relevance of cardiolipin-associated mitochondrial function during inflammation resolution. However, systemic SS-31 administration does not establish that the in vivo rescue is exclusively macrophage intrinsic. SS-31 may influence macrophage efferocytosis, thymocyte apoptosis, or both, and previous studies have reported effects of SS-31 on apoptotic pathways in other settings ^68,69^. However, our in vitro findings provide a distinction between these possibilities. Using Annexin V-binding apoptosis assay in Jurkat T cells, we found that SS-31 does not interfere with staurosporine-and dexamethasone-induced apoptosis (Figure S3). Nevertheless, we cannot exclude effects in other cell types or under different experimental conditions. Consequently, macrophage-specific manipulation of cardiolipin metabolism and direct analysis of engulfment in additional inflammatory models will be necessary to determine the extent to which this pathway contributes to inflammation resolution in vivo. Recent evidence also supports the broader relevance of cardiolipin to immune-cell metabolic fitness, as cardiolipin contributes to regulatory T-cell function in experimental colitis ^67^.

Together, these findings identify reduced cardiolipin as a previously unrecognized feature of lipin-1-deficient macrophages and support cardiolipin homeostasis as an important component of the mitochondrial program that enables macrophage respiration and efferocytosis. Rather than establishing the biochemical mechanism through which lipin-1 controls cardiolipin, our findings define a functional connection between lipin-1, mitochondrial inner membrane homeostasis, cardiolipin abundance, and pro-resolving macrophage function. Determining how lipin-1 influences cardiolipin trafficking, synthesis, remodeling, or stability will be important for understanding how lipid organization of the mitochondrial inner membrane supports macrophage metabolic adaptation during inflammation resolution.

### Limitations of the study

Several limitations should be considered when interpreting these findings. First, bulk RNA sequencing does not resolve selective transcriptional programs or macrophage population heterogeneity and therefore cannot exclude more focused transcriptional effects of lipin-1. Second, our mitochondrial morphology analyses relied on static imaging and do not directly measure mitochondrial fission or fusion rates. Third, the mitochondrial fraction used for proteomic analysis was enriched rather than completely purified, and protein-abundance measurements do not directly establish changes in respiratory-complex assembly or activity. Fourth, although PTPMT1 depletion and SS-31 treatment provide complementary evidence linking cardiolipin-associated pathways to macrophage respiration and efferocytosis, neither intervention is uniquely specific to cardiolipin. The mechanism connecting lipin-1 to cardiolipin abundance also remains unresolved. In addition, systemic administration of SS-31 does not establish a macrophage-intrinsic rescue mechanism in vivo, and myeloid-specific lipin-1 deletion does not exclude contributions from other myeloid populations. Finally, most mechanistic experiments were performed in murine bone marrow-derived macrophages, and the relevance of the lipin-1–cardiolipin relationship to primary human macrophages and human inflammatory disease remains to be established.

## RESOURCE AVAILABILITY

### Lead contact

Further information and requests for resources and reagents should be directed to and will be fulfilled by the lead contact, Matthew Woolard

### Material availability

Reagents are available from the lead contact author upon a reasonable request.

## ACKNOWLEDGEMENT

This work was supported by the American Heart Association award 25PRE1373728 (to O.O.I.); the National Institutes of Health awards: GM134974 and HL163106 (to M.D.W.); the National Science Foundation award 2537597 and the following National Institutes of Health awards: HL182859, HL182756, HL167758, and HL180481 (to A.Y.J.); the National Institutes of Health grants: R01HL172970, R01HL145753, R01HL145753-01S1, R01HL145753-03S1, and R00 HL122354, LSUHSC-S CCDS

Grant-in-Aid, Chancellor’s high aim, and Pathway Research Award (to M.S.B.).

## AUTHOR CONTRIBUTIONS

Conceptualization: O.O.I., T.T.B., R.M.S., J.N-F., M.D.W.; Data Curation: O.O.I., C.R.P., J.N-F., T.T.B., R.M.S., M.D.W.; Formal Analysis: O.O.I., C.R.P., J.N-F., T.T.B., R.M.S., C.S.A.; Funding Acquisition: M.D.W., O.O.I., A.Y.J., M.S.B.; Investigation: O.O.I., C.R.P., J.N-F., T.T.B., R.M.S., Z.M., C.S.A., R.P., K.J.M., D.M., M.D.W.; Methodology: O.O.I., C.R.P., J.N-F., T.T.B., R.M.S., Z.M., C.S.A., R.P., K.J.M., D.M., M.S.B., A.Y.J., M.D.W.; Resources: O.O.I., R.P., L.K., A.A.; Supervision: M.D.W.; Validation: O.O.I., J.N-F., R.P., L.K.; Visualization: O.O.I., C.R.P., J.N-F., T.T.B., R.M.S., C.S.A., R.P., K.J.M., D.M., M.D.W.; Writing-Original draft: O.O.I., J.N-F., Z.M., M.D.W.; Project Administration: M.D.W.

## DECLARATION OF INTERESTS

The authors have declared that they have no conflicting interests.

## STAR METHODS

Detailed methods are provided in the online version of this paper and include the following.

- KEY RESOURCES TABLE
- EXPERIMENTAL MODEL AND SUBJECT DETAILS

`o` Animal models and ethical approval
`o` Dexamethasone-induced thymus injury model
- METHOD DETAILS

`o` Bone marrow-derived macrophages (BMDMs) generation
`o` Cell culture and experimental perturbations
`o` L-cell-conditioned medium generation
`o` Immortalized BMDMs (iBMDMs) generation
`o` PTPMT1-depleted iBMDMs generation
`o` siRNA transfection
`o` SS-31 treatment of BMDMs
`o` Bulk RNA sequencing and bioinformatic analysis
`o` Quantitative reverse-transcription PCR
`o` TOM-20 immunofluorescence and mitochondrial network analysis
`o` Mitochondrial DNA copy number assay
`o` Transmission electron microscopy (TEM) mitochondrial network analysis
`o` Immunoblotting (Western Blotting) assay
`o` Mitochondrial fraction enrichment
`o` Mass spectrometry LC-MS/MS proteomic analysis
`o` NAO fluorescence microscopy
`o` NAO flow cytometry
`o` Targeted cardiolipin lipidomics by LC-MS/MS
`o` Cardiolipin fluorometric assay
`o` Apoptotic cell generation and labeling
`o` In vitro efferocytosis assay
`o` Seahorse mitochondrial stress test
`o` Annexin V-Binding Apoptosis Assay
- QUANTIFICATION AND STATISTICAL ANALYSIS

## MATERIALS AND METHODS

### 1. Animal models and ethical approval

All mouse studies were approved by the LSU Health Shreveport Institutional Animal Care and Use Committee (P24-015). Mice were housed and cared for in accordance with National Institutes of Health guidelines. Myeloid-specific lipin-1 knockout mice (lipin-1mKO) were generated by crossing mice carrying LoxP sites flanking exon 7 of Lpin1 (C57BL/6J and SV129 background; provided by B.N.F.) ^70^ with C57BL/6J LysM-Cre mice from The Jackson Laboratory (B6.129P2-Lyz2^tm1(cre)Ifo/J^ (JAX #004781). Age-matched Lpin1 flox/flox littermates served as controls. Cre insertion was detected by genomic PCR using the following primers: AAGGAGGGACTTGGAGGATG (common forward); GTCACTCACTGCTCCCCTGT (wild-type reverse); and ACCGGTAATGCAGGCAAAT (mutant reverse). Unless otherwise specified for an individual experiment, 8-to 16-week-old male and female mice were used to generate bone marrow-derived macrophages (BMDMs). Mice were euthanized by isoflurane overdose (Dechra; 200-129), followed by cervical dislocation to ensure death.

### 2. Cell culture and common experimental conditions

Unless otherwise indicated, macrophages were stimulated with 40 ng/mL recombinant mouse IL-4 (Leinco Technologies; I-207) for 6 h. RNA-sequencing experiments used a 4-h IL-4 stimulation. Untreated cells received no IL-4 and served as baseline controls, where indicated.

### 3. Bone marrow-derived macrophage generation

Femurs were flushed with unsupplemented DMEM (Gibco; 11965-092). Bone marrow cells were centrifuged and subjected to red blood cell lysis using 1× ammonium chloride-potassium (ACK) lysis buffer for 3 min at 4°C. After ACK lysis was washed off, cells were passed through a 40-µm cell strainer. 1 × 10^^7^ cells from lipin-1mKO mice were incubated with 10 µg/mL Tat-Cre recombinase (Excellgen EG-1001) in unsupplemented DMEM for 1 h before differentiation, while cells from littermate controls were not treated with Tat-Cre. 1 × 10^7^ bone marrow cells were differentiated for 7 d at 37 °C and 5% CO2 in growth medium consisting of KnockOut DMEM (Gibco; 10829-018), 30% L-cell-conditioned medium, 20% FBS (Atlas Biologicals; FP-0500-A), 1 mM sodium pyruvate (Corning; 25-000-CI), 2 mM GlutaMAX (Gibco; 35050-061), 100 U/mL penicillin-streptomycin (Gibco; 15140-122), and 0.2% sodium bicarbonate (Gibco; 25080-094) in 600-mL untreated tissue culture flasks. Differentiation medium was replaced on day 5. BMDM identity was verified by morphological assessment using phase-contrast microscopy. At approximately 80% confluency, BMDMs were washed with calcium-and magnesium-free DPBS and detached using 11 mM EDTA (pH 7.6). BMDMs were maintained in D10 medium consisting of DMEM, 10% FBS, 2 mM GlutaMAX, 100 U/mL penicillin-streptomycin, and 1 mM sodium pyruvate.

#### L-cell-conditioned medium

L929 cells (ATCC; CCL-1) were cultured in RPMI 1640 (Gibco; 11875-093) containing 10% FBS, 2 mM GlutaMAX, 1 mM sodium pyruvate, and 100 U/mL penicillin-streptomycin. Cells (3.75 × 10^5^) were seeded in T225 flasks containing 75 mL medium and cultured for 12–14 d at 37 °C until confluent, with a cobblestone appearance. Conditioned medium was collected, centrifuged at 500 RCF for 10 min at 4 °C, filtered through a 0.2-µm filter, stored at 4 °C for 24 h and −20 °C for 24 h, and then stored at −80 °C.

#### Derivation of immortalized BMDMs

Control and Lipin-1-deficient BMDMs were immortalized with the J2 murine retrovirus. J2-producing cells were a kind gift from Dr. Jonathan Kagan (Harvard Medical School) ^71^. The J2 virus-containing complete DMEM medium was produced by collecting the culture medium from J2-producing cells that were cultured for 24 hours after reaching 100% confluency. The virus-containing medium was centrifuged (400 x *g*, 5 min), and the supernatant was passed through a 0.2µm filter. BMDMs were transduced with J2 virus-containing supernatant fortified with 30% L-929 conditioned medium on consecutive days. Twelve hours after the second round of transduction, the medium was replaced with complete DMEM containing L-929 conditioned medium (30% v/v). To complete the immortalization process, BMDMs were split weekly (1:2 or 1:3) and the amount of L-929 conditioned medium was progressively reduced after every split (from 30% → 25% → 20% → 15% → 10% → 5% → 4% → 3% → 2% → 1% → 0%). Once weaned from L-929-conditioned medium, iBMDMs were cultured in complete DMEM medium. After cells were weaned, lipin-1-deficient iBMDMs were single-cell sorted and allowed to grow out. Samples were taken and lysed for lipin-1 quantification by immunoblotting. We identified 10 lipin-1-deficient clones; these 10 clones were mixed to create our lipin-1-deficient iBMDMs.

#### Generation of PTMTP1-depleted iBMDMs

Doxycycline-inducible shRNA-expressing macrophage cell lines were generated using lentiviral vectors targeting *Ptpmt1*. The constructs used in combination to deplete *Ptpmt1* were pLV[shRNA]-TerR:IRES:Puro-U6/2xTetO>mPtpmt1 [shRNA#1] (Vector Builder; LVI(VB251107-1310kmx)) and pLV[shRNA]-TetR:IRES/2xTetO>mPtpmt1[shRNA#2] (Vector Builder; LVI(VB251107-1313hgc)). Control shRNA construct was pLV[shRNA]-TerR:IRES:Puro-U6/2xTetO>Scramble_shRNA (Vector Builder; LVC(VB010000-9913att)-b). Each vector contains a tetracycline-responsive shRNA expression cassette and a TetR:IRES cassette to permit antibiotic selection of successfully transduced cells.

Macrophages were transduced with the lentiviral particles and subsequently cultured in medium containing puromycin (3µg/mL) to select for cells containing the integrated lentiviral construct. Following the establishment of puromycin-resistant populations, cells were maintained under standard culture conditions. To induce expression of the target-specific shRNA, cells were treated with 1µg/mL doxycycline for 72 h before experimental analysis. Parallel cells maintained in the absence of doxycycline served as non-induced controls. Knockdown efficiency was validated by immunoblot analysis, and depletion of PTPMT1 protein was confirmed before using the cells in downstream experiments.

### 3. Experimental perturbations siRNA transfection

WT and lipin-1mKO BMDMs (7.5 × 10^5^ cells) were seeded in T25 flasks in antibiotic-free growth medium. Lipofectamine RNAiMAX-siRNA complexes were prepared using 15 µL RNAiMAX (Invitrogen; 31985070) diluted in 500 µL Opti-MEM (Gibco; 31985070) and combined with 200 nM non-targeting siRNA (Horizon Discovery; D-001810-01-50), Drp1 siRNA (Qiagen; SI00982226), Opa1 siRNA (Qiagen; SI01365707), or Ptpmt1 siRNA (GeneSolution; 1027416). Cells were incubated with the transfection complexes overnight. The medium was then replaced with antibiotic-free growth medium, and cells were incubated in 5 mL of culture medium for a total of 72 h before downstream assays. Knockdown efficiency was confirmed by immunoblotting.

#### SS-31 treatment of BMDMs

SS-31 (Selleckchem; S9803) was prepared in water. Drug concentrations were optimized independently for each assay. For efferocytosis and cardiolipin-targeted LC-MS/MS experiments, lipin-1mKO BMDMs were pretreated with 0.5 µM SS-31 for 2 h and then stimulated with 40 ng/mL IL-4 for an additional 6 h in the continued presence of SS-31. Vehicle-treated lipin-1mKO and WT BMDMs received an equivalent volume of water (vehicle) in assay medium. For Seahorse XF Cell Mito Stress Test experiments, lipin-1mKO BMDMs were pretreated with 10 µM SS-31 2 hours before IL-4 stimulation. The higher concentration was selected based on assay-specific optimization to achieve reproducible effects on mitochondrial respiratory parameters.

### 4. Gene-expression analyses

#### Bulk RNA sequencing and bioinformatic analysis

BMDMs were generated from 8-to 12-week-old female lipin-1mKO mice and littermate controls. Genotypes were determined by genomic PCR, and reduced Lpin1 expression in lipin-1mKO BMDMs was confirmed by qRT-PCR. BMDMs were plated at 5 × 10^5^ cells per well in 24-well plates and allowed to adhere overnight. BMDMs from each mouse were divided into matched untreated and IL-4-treated cultures. Treated cultures received 40 ng/mL recombinant mouse IL-4 for 4 h.

Total RNA was isolated using the RNeasy Plus Mini Kit (Qiagen; 74134), including genomic DNA removal with the supplied gDNA Eliminator columns. RNA concentration was measured using a Qubit RNA assay, and integrity was assessed using an Agilent 2200 TapeStation. RNA samples meeting core quality requirements were used for library preparation with 500 ng RNA per sample.

Strand-specific, poly(A)-selected libraries were prepared using the Illumina TruSeq Stranded mRNA Library Preparation Kit. Fragment-size distributions were evaluated using an Agilent 2200 TapeStation D1000 assay, and library concentrations were determined using the Quanta PerfeCTa NGS Library Quantification Kit. Libraries were normalized to 4 nM, pooled, denatured, and diluted to approximately 1.8 pM with 1% PhiX at 1.8 pM. Libraries were sequenced in a single run on an Illumina NextSeq 550 using paired-end 2 × 75-bp reads. Base calling, quality scoring, demultiplexing, and FASTQ generation were performed using Illumina software by the LSU Health Sciences Center Shreveport Research Core Facility (RRID: SCR_024775).

FASTQ files were consolidated by sample before alignment. Sequencing generated an average of 38.6 million paired-end fragments per library (range, 32.7–43.6 million). Reads were aligned to the Mus musculus GRCm38.p6/mm10 reference genome using STAR 2.5.3a_modified and GENCODE M23 annotations. Alignment used two-pass basic mode (--twopassMode Basic), transcriptome-aligned output (--quantMode TranscriptomeSAM), and uniquely mapping reads (--outFilterMultimapNmax 1); all other parameters were maintained at default values. Unique mapping averaged 91.0% (range, 90.5–91.6%). Gene-level expected counts and transcripts per million (TPM) values were calculated using RSEM 1.2.31.

The RSEM expected-count matrix was analyzed in iDEP (version 2.4.4) using its default count-based differential-expression workflow. The primary comparison was IL-4-treated WT versus IL-4-treated lipin-1mKO BMDMs. No external TPM-based filter was applied. Differentially expressed genes were defined using a Benjamini-Hochberg-adjusted false-discovery rate of 0.10 and an absolute log2 fold-change threshold of 0.263, corresponding to a minimum 1.2-fold difference. Analysis was performed by the CAIPP Bioinformatics and Modeling Core (RRID: SCR_024776)

Gene Ontology Biological Process overrepresentation analysis was performed in iDEP on upregulated and downregulated genes separately using default pathway-size, multiple-testing, and background-gene settings. Terms with adjusted P < 0.05 were considered enriched. Transcription-factor target enrichment was performed using Enrichr and the ENCODE TF ChIP-seq library, with enrichment evaluated using adjusted P values. The final iDEP and Enrichr analyses were performed in June 2026.

Principal-component analysis was performed in GraphPad Prism 11.0.0 using untransformed TPM values after removing genes with zero TPM across all libraries. Volcano plots were generated using log2 fold change and −log10 Benjamini-Hochberg-adjusted P values.

#### Quantitative reverse-transcription PCR

WT and lipin-1mKO BMDMs (5 × 10^6^ cells) were plated in 12-well plates, allowed to adhere overnight, and treated with 40 ng/mL recombinant mouse IL-4 for 6 h. Total RNA was extracted using the GeneJET RNA Purification Kit (Thermo Fisher Scientific; K0731). Following the manufacturer’s instructions, 20 ng RNA was used for cDNA synthesis with the Verso cDNA Synthesis Kit (Thermo Fisher Scientific; AB-1453/B). Quantitative PCR was performed on a CFX Opus 96 Real-Time PCR System (Bio-Rad; 12011319) using SYBR Green SuperMix, ROX (Quantabio; 95055-500). Each reaction contained 5 µM forward and reverse primers. No-template and no-reverse-transcriptase controls were included. Cycling consisted of an initial denaturation at 95 °C for 3 min, followed by 39 cycles of 95 °C for 10 s and 55 °C for 30 s, with fluorescence acquisition at the end of each extension step. A melt-curve analysis was performed from 65 °C to 95 °C in 0.5 °C increments every 5 s to verify amplicon specificity. Verified primer sequences were obtained from the Harvard PrimerBank database. Expression of *Acadl*, *Echs1*, *Hadha*, *Acat1*, *Sptlc1*, *Degs1, Cers2*, *Cds1*, *Pgs1*, *Ptpmt1*, and *Crls1* were measured. Relative expression was calculated by the 2^−ΔΔCt^ method after normalization to Gapdh and expressed relative to untreated WT BMDMs. All primer sequences were obtained from the Harvard Primer Bank and were validated and tested for efficiency.

**Table 1.**
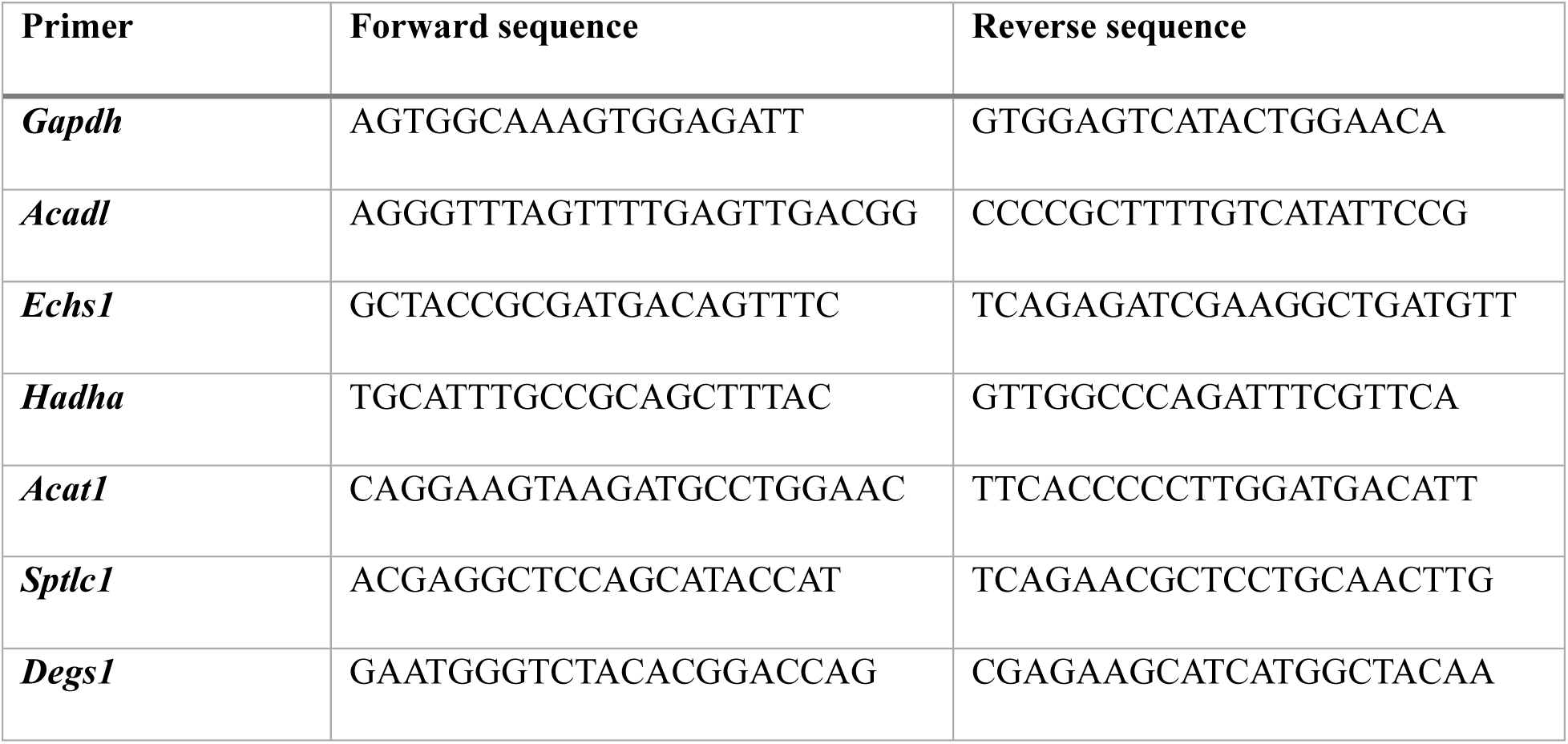
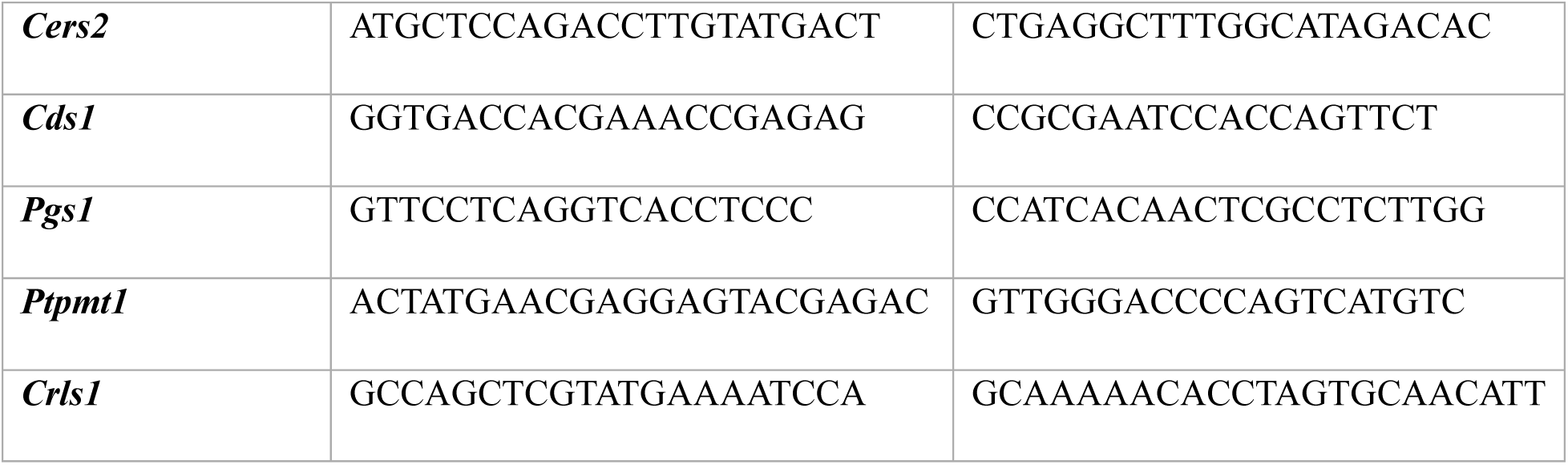
Quantitative PCR primer sequences.

### 5. Mitochondrial morphology and ultrastructure

#### TOM-20 immunofluorescence and mitochondrial-network analysis

BMDMs (1.75 × 10^5^ cells) were plated on 12-mm glass coverslips in 24-well plates and treated with or without IL-4 for 6 h. Cells were fixed in 4% paraformaldehyde (Electron Microscopy Sciences; 15714-S) for 30 min at room temperature, quenched with 0.1 M glycine for 30 min, and permeabilized and blocked for 1 h in PBS containing 1% BSA, 0.1% cold-water fish-skin gelatin, 0.1% Tween 20, and 0.05% sodium azide. Cells were incubated with TOM20 antibody (Abcam; AB56783; 1:250) for 1 h, and Alexa Fluor 568 donkey anti-mouse IgG (Thermo Fisher Scientific; A10037; 1:200) for 90 min at room temperature. Nuclei were stained with DAPI (Invitrogen; D3571; 1:36,400) for 15 min. Coverslips were mounted using VECTASHIELD HardSet Antifade Mounting Medium (Vector Laboratories; H-1400). Images were acquired using a Nikon SIM-E/A1-R confocal system with a 60× objective, no digital zoom, line averaging of 2, and 1024 × 1024-pixel acquisition at a single optical section. Raw images were deconvolved using a custom convolution filter in ImageJ. Deconvolved images were analyzed in NIS-Elements using the object-count tool with restrictions based on length, width, circularity, and area. Mitochondrial length and circularity were summarized for each field relative to the number of DAPI-stained nuclei. All mitochondrial analyses were performed blinded to genotype and treatment.

#### Mitochondrial DNA copy-number assay

BMDMs (1 × 10^6^ cells) were treated with or without 40 ng/mL IL-4 for 6 h. Genomic DNA was isolated using the GeneJET Genomic DNA Purification Mini Kit (Thermo Fisher Scientific; K0781), quantified using a NanoDrop spectrophotometer, and 2 ng DNA was used per qPCR reaction. Real-time PCR was performed on a CFX Opus 96 Real-Time PCR System (Bio-Rad; 12011319) using SYBR Green SuperMix, ROX (Quantabio; 95055-500), and the NovaQUANT Mouse Mitochondrial-to-Nuclear DNA Ratio Kit (MilliporeSigma; 72621-1KIT), which amplifies two nuclear genes (BECN1 and NEB) and two mitochondrial genes (trLEV and 12s). Cycling consisted of an initial denaturation at 95 °C for 10 min, followed by 40 cycles of 95 °C for 3 s and 60 °C for 30 s, with fluorescence acquisition at the end of each extension step. A melt-curve analysis was performed from 65 °C to 95 °C in 0.5 °C increments every 5 s. Relative mitochondrial DNA copy number was calculated for each mitochondrial-to-nuclear gene pair (trLEV/BECN1 and 12s/NEB) as 2^ΔCt^, where ΔCt = Ct_nuclear − Ct_mitochondrial. The final value was reported as the mean of the two mitochondrial-to-nuclear ratios.

#### Transmission electron microscopy mitochondrial analysis

BMDMs were treated with or without IL-4 for 6 h and fixed in a mixture of 2.5% glutaraldehyde and 2% formaldehyde in 0.1M Sorensen’s buffer, pH 7.4. Post fixation was done using reduced osmium textroxide (1% osmium tetroxide/1% K3FeCN6 in 0.1M Sorensen’s buffer pH 7.4) followed by 0.5% uranyl acetate aqueous en bloc stain (overnight) at 4°C. The cells were dehydrated with a graded series of ethanols, followed by propylene oxide and infiltrated with Durcupan Araldite resin (Sigma) according to standard protocol ^72^. Samples were processed and imaged by the LSU Health Sciences Center Shreveport Electron Microscopy Core (RRID: SCR_024775). Images with mitochondria in focus were selected across groups for quantification. ImageJ was used to quantify mitochondrial numbers and morphology. Images containing entire cell profiles were analyzed using the Multi-Point tool to enumerate mitochondrial profiles. Individual mitochondrial profiles were manually traced using the Freehand Selection tool, and morphometric measurements, including area, roundness, and circularity, were obtained using the Set Measurements and Measure functions.

#### 6. Protein and mitochondrial fraction analyses Cell lysis and immunoblotting

Cells were lysed in 1× NuPAGE lysis buffer supplemented with 100 mM DL-dithiothreitol (Sigma-Aldrich; D0632-10G), 1× phosphatase inhibitor cocktail 2 (Sigma-Aldrich; P5726-5ML), 1× phosphatase inhibitor cocktail 3 (Sigma-Aldrich; P0044-5ML), and 1× Halt protease inhibitor (Thermo Fisher Scientific; 181279). Cell lysates were sonicated and heat-denatured, and protein concentration was determined using the Pierce 660 nm Protein Assay (Thermo Fisher Scientific; 22660). Equal amounts of protein (20 µg) were loaded on NuPAGE 4–12% Bis-Tris Midi Gels (Thermo Fisher Scientific; WG1401BOX) and separated in 1× MOPS or 1× MES running buffer at 200 V and 400 mA for 53 min. Proteins were transferred to Immobilon-P PVDF membranes (Sigma-Aldrich; SIPVH00010) at 20 V and 400 mA for 45 min. Membranes were blocked with Intercept (TBS) Blocking Buffer (LI-COR Biosciences; 927-60001) for 1 h at room temperature and incubated with primary antibodies overnight at 4 °C. After three 15-min washes in 1× Tris-buffered saline containing Tween-20 (TBST), membranes were incubated with species-specific HRP-conjugated secondary antibodies diluted in 5% nonfat dry milk (OmniBlok; AmericanBio; AB10109-01000) containing 0.01% sodium dodecyl sulfate for 2 h at room temperature. Membranes were washed three times in TBST, and protein bands were visualized using WesternBright ECL Spray (Advansta; K-12049-D50).

#### Mitochondrial Isolation Buffer

Mitochondrial isolation buffer (MIB) consisted of 0.25 M sucrose, 5 mM HEPES, and 3.1 mM EGTA dissolved in distilled water to a final volume of 10 mL ^73^. For buffer preparations containing BSA, fatty acid–free BSA was added to a final concentration of 1% (w/v). MIB was stored at −20°C for up to 3 months before replacement.

#### Mitochondrial Isolation

BMDMs were cultured for 7–10 days and resuspended in mitochondrial isolation buffer (MIB) containing 1% BSA. Macrophages were transferred to custom 1.5 mL microcentrifuge tubes suitable for Dounce homogenization. The entire homogenate was centrifuged at 900 × g for 10 minutes at 4°C, and the resulting supernatant was transferred to fresh 1.5 mL microcentrifuge tubes while the pellet was retained.

Supernatants were further centrifuged at 10,000 × g for 10 minutes at 4°C. The resulting supernatants were discarded, and pellets were resuspended in 50 µl of MIB containing BSA, then centrifuged again at 10,000 × g for 10 minutes at 4°C. Supernatants were discarded, and mitochondrial pellets were resuspended in 50 µl MIB without BSA and centrifuged again at 10,000 × g for 10 minutes at 4°C. Following the final wash, cell pellets were lysed in 37.5 µl of 1X Radio Immuno-precipitation Assay (RIPA) buffer mixed with 12.5 µl of NuPAGE lysis buffer, and samples were stored at −80°C until analysis. Mitochondrial fractions were assessed for purity by checking TOM-20 and GAPDH intensity via immunoblot analysis. Whole-cell lysates collected at the time of harvesting were used as controls to establish baseline mitochondrial protein levels.

#### Mass Spectrometry and Proteomic Analysis

Protein (10 µg) from each sample was separated by SDS-PAGE and stained with Imperial Protein Stain (Thermo Scientific). Gel lanes were excised and cut into approximately 2 × 2 mm cubes, which were subjected to in-gel tryptic digestion, disulfide bond reduction with dithiothreitol, and cysteine alkylation with iodoacetamide using the Thermo Scientific™ Gel Tryptic Digestion Kit, according to the manufacturer’s protocol. Digested peptides were vacuum-dried and reconstituted in LC–MS–grade water containing 0.1% (v/v) formic acid for untargeted discovery proteomics analysis. LC-MS/MS analysis was performed on an Ultimate 3000 RSLCnano system coupled to an Orbitrap Exploris 480 mass spectrometer.

Reconstituted peptides (5.0 µl) were loaded onto a trap column (PepMap C18, 2 cm × 100 µm, 100 Å) at a flow rate of 20 µl/min in 0.1% formic acid and subsequently separated on an analytical column (EasySpray, 50 cm × 75 µm, C18, 1.9 µm, 100 Å) at 300 nl/min using a linear gradient from 5% to 45% solvent B (100% acetonitrile, 0.1% formic acid) over 120 minutes. Both precursor and fragment ions were acquired in the Orbitrap mass analyzer. Precursor ions were acquired over an m/z range of 375–1500 at a resolution of 120,000 (at m/z 200). Precursor fragmentation was performed using higher-energy collisional dissociation with a normalized collision energy (NCE) of 32, and fragment ions were acquired at a resolution of 150,000 (at m/z 200). Data were collected in a top-speed data-dependent acquisition mode with a 3-second cycle time between MS and MS/MS scans. Ion transfer capillary voltage was maintained at 2.1 kV. Raw mass spectrometry data were analyzed using Proteome Discoverer (version 3.2, Thermo Fisher Scientific) with SequestHT and a species-specific FASTA database, followed by Percolator peptide validation. Cysteine alkylation was specified as a fixed modification, and deamidation of asparagine residues was included as a variable modification.

#### Proteomic Statistical Analysis and Databases Used

Following initial mass spectrometry data processing, protein abundance data were further filtered and analyzed to identify genotype-and treatment-dependent changes in the mitochondrial proteome. The initial mass spectrometry dataset contained 5,443 protein-group identifications. To restrict subsequent analyses to bona fide mitochondrial proteins, identified proteins were cross-referenced with **MitoCarta 3.0**. Proteins not represented in MitoCarta 3.0 were excluded from mitochondrial-focused analyses. This filtering resulted in a dataset of **562 MitoCarta 3.0–annotated mitochondrial proteins**. MitoCarta 3.0 annotations were also used to assign sub-mitochondrial localization, including mitochondrial matrix, inner mitochondrial membrane, intermembrane space, and outer mitochondrial membrane.

To minimize the influence of proteins detected inconsistently across biological replicates, mitochondrial proteins were retained for genotype-level quantitative analysis when abundance values were detected in at least **7 of 9 samples for both WT and KO cells** across experimental conditions. Applying this criterion resulted in **436 mitochondrial proteins** eligible for quantitative genotype-level analysis. For proteins meeting this detection threshold, missing abundance values were assigned a value of 0.05 to permit statistical analysis.

Unpaired *t*-tests were used to compare protein abundance between untreated WT and KO cells, LPS-treated WT and KO cells, and IL-4–treated WT and KO cells. An additional analysis compared protein abundance across all WT and KO samples independent of treatment. Fold changes were calculated for each comparison and used together with statistical significance to identify proteins altered by loss of lipin-1. Proteins exhibiting changes in abundance were subsequently evaluated individually and at the pathway level.

Functional annotation and mitochondrial pathway assignments were obtained primarily from **MitoCarta 3.0**, with UniProt used as an additional resource to verify protein function and resolve annotations when necessary.

### 7. Cardiolipin analyses

#### NAO staining and flow cytometry

BMDMs were resuspended at 5 × 10^5^ cells in 0.5 mL D10 medium and treated with or without IL-4 (40 ng/mL) for 6 h at 37 °C. Cells were stained with 0.5 µM nonyl acridine orange (NAO; Biotium; 70012) in PBS containing 1% BSA for 30 min at 37 °C in the dark and fixed in 4% formaldehyde for 30 min at 4 °C. Data were acquired on a NovoCyte Quanteon using NovoExpress 1.6.2. NAO fluorescence was detected using the 488-nm laser and the FITC-equivalent detector. At least 50,000 events were collected per sample. Debris and doublets were excluded sequentially, and the macrophage gate was defined using unstained controls. NAO fluorescence was quantified as median fluorescence intensity within the macrophage gate. Median fluorescence intensity values were normalized to the untreated WT group for statistical analysis. Histograms show normalized event frequency.

#### NAO fluorescence microscopy

BMDMs (1.25 × 10^5^ cells) were plated on 12-mm coverslips in 24-well tissue-culture-treated plates and treated with IL-4 (40 ng/mL) for 6 h. Cells were fixed in 4% formaldehyde for 30 min at room temperature, incubated with 5 µM NAO (Biotium; 70012) for 15 min, followed by three PBS washes. Nuclei were stained with DAPI for 15 min, and coverslips were mounted using SlowFade Gold Antifade Mountant with DAPI (Invitrogen; S36938). Images were acquired at 20× magnification using a Keyence BZ-X810 microscope. NAO fluorescence was quantified in ImageJ as integrated density divided by the number of cells per image.

#### Targeted cardiolipin lipidomics

BMDMs (2 × 10^6^ cells) were seeded in 2 mL D10 medium in 15-mL tubes and treated with or without IL-4 (40 ng/mL) for 6 h. Cell pellets were stored at −80 °C until lipid extraction and analysis by the LSU Health Shreveport Research Core Facility (RRID: SCR_024775). Samples were thawed, and 400 µL of each macrophage sample was transferred to a 1.5-mL microcentrifuge tube. Methanol (500 µL) was added, samples were vortexed, followed by the addition of 250 µL chloroform. Samples were allowed to separate for 10 min, after which 350 µL of deionized water was added to facilitate phase separation. Samples were centrifuged at 12,000 rpm for 10 min, and 200 µL of the lower chloroform phase was transferred to a glass vial and evaporated to dryness in a fume hood. Dried lipid extracts were reconstituted in 100 µL acetonitrile.

LC–MS analysis was performed using a Thermo Fisher Vanquish Horizon UHPLC coupled to an Orbitrap Exploris 120 mass spectrometer. Lipids were separated on a Thermo Accucore C30 column (2.6 µm, 150 × 2.1 mm) maintained at 45 °C at a flow rate of 0.26 mL/min. Mobile phase A consisted of acetonitrile (60:40, v/v) containing 0.1% formic acid, and mobile phase B consisted of isopropyl alcohol (90:10, v/v) containing 0.1% formic acid. The chromatographic gradient was programmed as follows: 30% B at 0 min, 43% B at 2 min, 55% B at 2.1 min, 65% B at 12 min, 85% B at 18 min, 100% B from 20 to 25 min, and return to 30% B at 25.1 min, which was maintained through 31 min. A 5-µL aliquot of each sample was injected for analysis.

Ionization was performed using heated electrospray ionization (H-ESI) with static spray voltages of 3,500 V in positive-ion mode and 2,400 V in negative-ion mode. Source conditions included sheath gas at 50 arbitrary units, auxiliary gas at 10 arbitrary units, sweep gas at 1 arbitrary unit, and ion-transfer-tube and vaporizer temperatures of 300 °C. Cardiolipin species were identified based on their chromatographic and mass-spectrometric characteristics. Because absolute quantification using external calibration curves was not used for the final analysis, cardiolipin abundance was determined from the integrated LC–MS peak area for each identified species. Peak-area values were subsequently normalized to the mean value of the untreated WT group for each cardiolipin species and are presented as relative cardiolipin abundance.

#### Cardiolipin fluorometric assay

Cardiolipin levels were quantified using a Cardiolipin assay kit (Abcam; ab241036) according to the manufacturer’s instructions. Briefly, 1 x 10^6^ iBMDMs were seeded and stimulated with IL-4 for 6 h. Cells were harvested with ice-cold 11 mM EDTA, centrifuged, and the supernatant discarded. Cell pellets were resuspended in Assay Buffer and lysed by sonication. Protein concentration of cell lysates was determined using the Pierce 660 nm Protein Assay (Thermo Fisher Scientific; 22660). Equal amounts of protein (10 µg per sample) were incubated with the cardiolipin (CL) probe for 10 minutes, and fluorescence was measured at Excitation/Emission of 340/480nm. Cardiolipin concentrations were calculated using the Standard curve readings and were represented as nmol/mL.

### 8. Macrophage functional assays Apoptotic-cell generation and labeling

Jurkat cells (ATCC; TIB-152) were maintained in RPMI 1640 supplemented with 10% FBS, 2 mM GlutaMAX, 10 mM HEPES, and 100 U/mL penicillin-streptomycin. To induce apoptosis, cells were washed, resuspended in PBS at 1–1.5 × 10^6^ cells/mL, and exposed to UV irradiation for 10 min in 100-mm Petri dishes using a UV Clave Ultraviolet Chamber (Benchmark Scientific; B1450). Cells were collected, pelleted at 500 RCF rpm for 5 min, and labeled with 5 µM CFSE (BioLegend; 423801) in serum-free DMEM at 10–100 × 10^6^ cells/mL for 20 min at room temperature. Labeling was quenched with DMEM containing 10% FBS at five times the labeling volume. Cells were washed, resuspended in serum-free DMEM, and incubated for 2 h at 37 °C before use in efferocytosis assays. Annexin V staining confirmed that >80% of cells were apoptotic.

#### In vitro efferocytosis assay

siRNA-transfected BMDMs (1.75 × 10^5^ cells) were plated on 12-mm coverslips in 24-well tissue-culture-treated plates and treated with IL-4 (40 ng/mL) for 6 h. BMDMs were incubated with CFSE-labeled apoptotic cells at a 1:4 macrophage-to-apoptotic-cell ratio for 45 min. Unengulfed apoptotic cells were removed by three PBS washes, and BMDMs were fixed in 4% paraformaldehyde for 30 min. Nuclei were stained with DAPI for 15 min, and coverslips were mounted using SlowFade Gold Antifade Mountant with DAPI (Invitrogen; S36938). Images were acquired at 20× magnification using a Keyence BZ-X810 microscope. Macrophages and engulfed apoptotic cells were counted using the ImageJ 1.54g Multi-Point tool. Efferocytosis was quantified as the percentage of macrophages containing apoptotic cells: (number of macrophages containing apoptotic cells / total number of macrophages) × 100. For SS-31 experiments, lipin-1mKO BMDMs were pretreated with 0.5µM SS-31 for 2 h and then stimulated with 40 ng/mL IL-4 for an additional 6 h in the continued presence of SS-31; WT and lipin-1mKO vehicle-control cells received water vehicle.

#### Seahorse mitochondrial stress test

BMDMs (1 × 10^5^ cells) were seeded in Seahorse XF24 V7 PS Cell Culture Microplates (Agilent Technologies; 100777-004) and subjected to the indicated treatments. Following treatment, cells were washed twice and incubated in Seahorse XF DMEM Medium, pH 7.4 (Agilent; 103575-100), supplemented with 2 mM GlutaMAX (Gibco; 35050-061), 10 mM glucose (Agilent; 103577-100), and 1 mM pyruvate (Agilent; 103578-100). Cells were equilibrated for 1 h at 37 °C in a non-CO2 incubator before analysis. Oxygen-consumption rate (OCR) was measured using an Agilent Seahorse XFe24 Analyzer. The Mito Stress Test used sequential injections of 1 µM oligomycin (Sigma-Aldrich; 75351-5MG), 2 µM FCCP (Sigma-Aldrich; SML2959-1ML), and rotenone plus antimycin A (Sigma-Aldrich; R8875-1G and A8674-25MG). For SS-31 experiments, lipin-1mKO BMDMs were pretreated with 10µM SS-31 for 2 h and then stimulated with IL-4 for an additional 6 h in the continued presence of SS-31; WT and lipin-1mKO vehicle-control cells received water vehicle.

### 9. Dexamethasone-induced in vivo thymus efferocytosis Animal treatment and tissue collection

Lipin-1mKO mice received a single intraperitoneal injection of SS-31 (5 mg/kg; Selleckchem; S9803) or vehicle, and WT littermate controls received vehicle. SS-31 or water vehicle was diluted in PBS for injection. Six hours later, all mice received 250 µg dexamethasone (Sigma-Aldrich; D902-25MG) in 250 µL PBS by intraperitoneal injection. Mice were euthanized 18 h after dexamethasone administration by isoflurane overdose, and thymi were harvested and weighed. One thymic lobe was processed for single-cell analysis, and the contralateral lobe was fixed in 10% neutral-buffered formalin and paraffin-embedded for histologic analysis.

#### Thymic single-cell preparation and flow cytometry

One thymic lobe from each mouse was mechanically dissociated to generate a single-cell suspension. Red blood cells were lysed in 1× ACK lysis buffer for 5 min, and total cell numbers were determined. Cells were stained with ApoTracker Green (BioLegend; 750000920) to identify apoptotic cells, PE/Cyanine7 anti-mouse/human CD11b antibody (BioLegend; 101215), and Brilliant Violet 650 anti-mouse F4/80 antibody (BioLegend; 123149) to identify the macrophage population. Flow cytometry was used to quantify apoptotic-cell accumulation and macrophage abundance.

#### TUNEL/Mac-2 histology and in situ efferocytosis analysis

The contralateral thymic lobe was fixed in 10% neutral-buffered formalin, paraffin-embedded, sectioned, and stained using a TUNEL apoptosis kit (Biotium; 30074) together with anti-Mac-2 antibody (BioLegend; 125410). Images were acquired at 20× magnification using a Keyence BZ-X810 microscope. TUNEL-positive apoptotic cells were classified as either associated with Mac-2-positive macrophages or free within the tissue. In situ efferocytosis was expressed as the ratio of macrophage-associated TUNEL-positive cells to free TUNEL-positive cells.

#### Annexin V-Binding Apoptosis Assay

Jurkats were treated with 0.5 µM or 10 µM SS-31 or vehicle for 2 hours before no treatment or treatment with 2.5 µM staurosporine (MedChemExpress; HY-15141) or 500 µM dexamethasone (Sigma Aldrich; D902-25MG) for additional 4 hours to induce apoptosis. Apoptosis was measured by Annexin V-binding assay by adding one drop of Annexin V Ready Flow Reagent (Thermo Fisher Scientific; R37175) to 1 x

10^5^ cells suspended in 500 µL of homemade annexin V-binding buffer followed by incubation for 15 minutes at room temperature. Cells were counted in Countess 3 FL automated cell counter.

### 10. Statistical analysis

The figure legends report the number and type of replicates and indicate whether individual data points represent biological or technical replicates. Technical replicates were averaged to produce one value per biological replicate before statistical analysis unless otherwise indicated. Statistical analyses were performed using GraphPad Prism 11.0.0 unless otherwise specified. The statistical test, summary measure, and significance notation for each analysis are reported in the corresponding figure legend. Tests were two-sided, and *p* < 0.05 was considered significant unless a method-specific adjusted threshold is stated.

For the mitochondrial proteomics compartment analysis, the association between sub-mitochondrial localization (matrix, MIM, or MOM) and directional abundance category (decreased, unchanged, or increased using the ±10% operational threshold described above) was evaluated using a Pearson chi-square test of independence, and Cramér’s V was calculated as an effect-size estimate. Pairwise comparisons of the proportion of decreased versus not-decreased proteins between compartments were performed using two-sided Fisher’s exact tests with Holm correction. Fold-change distributions were also analyzed without categorization using a Kruskal-Wallis test followed by pairwise two-sided Mann-Whitney U tests with Holm correction.

## Notes

### Competing Interest Statement

The authors have declared no competing interest.

## REFERENCES

1. Aksentijevich, M., Lateef, S. S., Anzenberg, P., Dey, A. K. & Mehta, N. N. Chronic Inflammation, Cardiometabolic Diseases and Effects of Treatment: Psoriasis as a Human Model. Trends Cardiovasc Med 30, 472–478 (2020).

2. Saas, P., Vetter, M., Maraux, M., Bonnefoy, F. & Perruche, S. Resolution therapy: Harnessing efferocytic macrophages to trigger the resolution of inflammation. Front Immunol 13, 1021413 (2022).

3. Zhang, S. et al. Efferocytosis Fuels Requirements of Fatty Acid Oxidation and the Electron Transport Chain to Polarize Macrophages for Tissue Repair. Cell Metabolism 29, 443–456.e5 (2019).

4. Schilperoort, M. et al. The role of efferocytosis-fueled macrophage metabolism in the resolution of inflammation. Immunol Rev 319, 65–80 (2023).

5. Gao, J. et al. Macrophage efferocytosis promotes inflammation resolution and accelerates wound healing. Commun Biol 9, 613 (2026).

6. Barnig, C., et al. Resolution of Inflammation after Skeletal Muscle Ischemia– Reperfusion Injury: A Focus on the Lipid Mediators Lipoxins, Resolvins, Protectins and Maresins. Antioxidants 11, (2022).

7. Schilke, R. M. et al. Macrophage-Associated Lipin-1 Promotes β-Oxidation in Response to Proresolving Stimuli. Immunohorizons 4, 659–669 (2020).

8. Bamgbose, T. T. et al. Lipin-1 restrains macrophage lipid synthesis to promote inflammation resolution. The Journal of Immunology 214, 85–103 (2025).

9. Blackburn, C. M. R. et al. Myeloid-associated lipin-1 transcriptional co-regulatory activity is atheroprotective. Atherosclerosis 330, 76–84 (2021).

10. Bradshaw, P. C. et al. The Role of Cardiolipin in Brain Bioenergetics, Neuroinflammation, and Neurodegeneration. Mol Neurobiol 62, 7022–7040 (2025).

11. Chai, Z. et al. Cardiolipin’s multifaceted role in immune response: a focus on interacting proteins. Front. Immunol. 16, (2025).

12. Jiang, Z. et al. Cardiolipin Regulates Mitochondrial Ultrastructure and Function in Mammalian Cells. Genes 13, (2022).

13. Romanova, N. et al. Cadmium-cardiolipin disruption of respirasome assembly and redox balance through mitochondrial membrane rigidification. Journal of Lipid Research 66, 100750 (2025).

14. Balasubramanian, K. et al. Dichotomous roles for externalized cardiolipin in extracellular signaling: Promotion of phagocytosis and attenuation of innate immunity. Sci Signal 8, ra95 (2015).

15. Coats, S. R. et al. Cardiolipins Act as a Selective Barrier to Toll-Like Receptor 4 Activation in the Intestine. Appl Environ Microbiol 82, 4264–4278 (2016).

16. Pizzuto, M. et al. Saturation of acyl chains converts cardiolipin from an antagonist to an activator of Toll-like receptor-4. Cell. Mol. Life Sci. 76, 3667–3678 (2019).

17. Wenzel, T. J., Ranger, A. L., McRae, S. A. & Klegeris, A. Extracellular cardiolipin modulates microglial phagocytosis and cytokine secretion in a toll-like receptor (TLR) 4-dependent manner. Journal of Neuroimmunology 353, 577496 (2021).

18. Reynolds, M. B. et al. Cardiolipin coordinates inflammatory metabolic reprogramming through regulation of Complex II disassembly and degradation. Science Advances 9, eade8701 (2023).

19. Iyer, S. S. et al. Mitochondrial Cardiolipin Is Required for Nlrp3 Inflammasome Activation. Immunity 39, 311–323 (2013).

20. Chambers, K. T., et al. Myocardial Lipin 1 knockout in mice approximates cardiac effects of human *LPIN1* mutations. JCI Insight 6, (2021).

21. Finck, B. N. et al. Lipin 1 is an inducible amplifier of the hepatic PGC-1α/PPARα regulatory pathway. Cell Metabolism 4, 199–210 (2006).

22. Peterson, T. R. et al. mTOR Complex 1 Regulates Lipin 1 Localization to Control the SREBP Pathway. Cell 146, 408–420 (2011).

23. Wang, J. et al. Lipin1-dependent transcriptional inactivation of SREBPs contributes to selinexor sensitivity in multiple myeloma. Acta Pharmacol Sin 46, 2496–2508 (2025).

24. Benador, I. Y. et al. Mitochondria Bound to Lipid Droplets Have Unique Bioenergetics, Composition, and Dynamics that Support Lipid Droplet Expansion. Cell Metabolism 27, 869–885.e6 (2018).

25. Park, S. et al. Defective mitochondrial fission augments NLRP3 inflammasome activation. Sci Rep 5, 15489 (2015).

26. Wang, Y. et al. Mitochondrial Fission Promotes the Continued Clearance of Apoptotic Cells by Macrophages. Cell 171, 331–345.e22 (2017).

27. Susser, L. I. et al. Mitochondrial Fragmentation Promotes Inflammation Resolution Responses in Macrophages via Histone Lactylation. Mol Cell Biol 43, 531–546 (2023).

28. Huang, H. et al. piRNA-associated germline nuage formation and spermatogenesis require MitoPLD profusogenic mitochondrial-surface lipid signaling. Dev Cell 20, 376–387 (2011).

29. Lam, J. et al. A Universal Approach to Analyzing Transmission Electron Microscopy with ImageJ. Cells 10, 2177 (2021).

30. Malachowski, A. et al. Mitochondrial dynamics and antiviral responses in Atlantic salmon cardiomyocytes during piscine myocarditis virus infection. iScience 29, 116831 (2026).

31. Ngo, J. et al. Mitochondrial morphology controls fatty acid utilization by changing CPT1 sensitivity to malonyl-CoA. EMBO J 42, EMBJ2022111901 (2023).

32. Trumpff, C. et al. Mitochondrial respiratory chain protein co-regulation in the human brain. Heliyon 8, e09353 (2022).

33. Cogliati, S. et al. Mitochondrial Cristae Shape Determines Respiratory Chain Supercomplexes Assembly and Respiratory Efficiency. Cell 155, 160–171 (2013).

34. Yoo, Y., Yeon, M., Yoon, M.-S. & Seo, Y.-K. Role of cardiolipin in skeletal muscle function and its therapeutic implications. Cell Commun Signal 23, 36 (2025).

35. Zhang, M., Mileykovskaya, E. & Dowhan, W. Gluing the Respiratory Chain Together: CARDIOLIPIN IS REQUIRED FOR SUPERCOMPLEX FORMATION IN THE INNER MITOCHONDRIAL MEMBRANE*. Journal of Biological Chemistry 277, 43553–43556 (2002).

36. Mileykovskaya, E. & Dowhan, W. Cardiolipin-dependent formation of mitochondrial respiratory supercomplexes. Chemistry and Physics of Lipids 179, 42–48 (2014).

37. Spera, I. et al. The J2-Immortalized Murine Macrophage Cell Line Displays Phenotypical and Metabolic Features of Primary BMDMs in Their M1 and M2 Polarization State. Cancers (Basel) 13, 5478 (2021).

38. Liu, G.-Y. et al. The phospholipase iPLA2γ is a major mediator releasing oxidized aliphatic chains from cardiolipin, integrating mitochondrial bioenergetics and signaling. Journal of Biological Chemistry 292, 10672–10684 (2017).

39. Szeto, H. H. First-in-class cardiolipin-protective compound as a therapeutic agent to restore mitochondrial bioenergetics. Br J Pharmacol 171, 2029–2050 (2014).

40. Chandran, S. et al. Lipin-1 Contributes to IL-4 Mediated Macrophage Polarization. Front Immunol 11, 787 (2020).

41. Luo, P. et al. SS31 alleviates LPS-induced acute lung injury by inhibiting inflammatory responses through the S100A8/NLRP3/GSDMD signaling pathway. Eur J Med Res 29, 567 (2024).

42. Yurdagul, A. et al. Macrophage Metabolism of Apoptotic Cell-Derived Arginine Promotes Continual Efferocytosis and Resolution of Injury. Cell Metab 31, 518–533.e10 (2020).

43. Gerlach, B. D. et al. Efferocytosis induces macrophage proliferation to help resolve tissue injury. Cell Metabolism 33, 2445–2463.e8 (2021).

44. Sun, C. et al. IRF-1 Regulates Mitochondrial Respiration and Intrinsic Apoptosis Under Metabolic Stress through ATP Synthase Ancillary Factor TMEM70. Inffammation 48, 2548–2562 (2025).

45. Wang, L. et al. The multiple roles of interferon regulatory factor family in health and disease. Sig Transduct Target Ther 9, 282 (2024).

46. Zong, Q. et al. Hypomethylated interferon regulatory factor 8 recruits activating protein-2α to attenuate porcine epidemic diarrhea virus infection in porcine jejunum. Front Immunol 14, 1187144 (2023).

47. Yang, D. et al. IFN Regulatory Factor 8 Sensitizes Soft Tissue Sarcoma Cells to Death Receptor-initiated Apoptosis via Repression of FLICE-like Protein Expression. Cancer Res 69, 1080–1088 (2009).

48. Tong, M., Zablocki, D. & Sadoshima, J. The role of Drp1 in mitophagy and cell death in the heart. J Mol Cell Cardiol 142, 138–145 (2020).

49. Fan, R. Z. et al. A partial Drp1 knockout improves autophagy flux independent of mitochondrial function. Mol Neurodegener 19, 26 (2024).

50. Quintana-Cabrera, R. et al. Opa1 relies on cristae preservation and ATP synthase to curtail reactive oxygen species accumulation in mitochondria. Redox Biology 41, 101944 (2021).

51. Frezza, C. et al. OPA1 Controls Apoptotic Cristae Remodeling Independently from Mitochondrial Fusion. Cell 126, 177–189 (2006).

52. Milani, M. et al. DRP-1 is required for BH3 mimetic-mediated mitochondrial fragmentation and apoptosis. Cell Death Dis 8, e2552 (2017).

53. Hall, A. R., Burke, N., Dongworth, R. K. & Hausenloy, D. J. Mitochondrial fusion and fission proteins: novel therapeutic targets for combating cardiovascular disease. Br J Pharmacol 171, 1890–1906 (2014).

54. Moltedo, O., Remondelli, P. & Amodio, G. The Mitochondria–Endoplasmic Reticulum Contacts and Their Critical Role in Aging and Age-Associated Diseases. Front Cell Dev Biol 7, 172 (2019).

55. Hinton, A. et al. ATF4 Dependent Increase in Mitochondrial-Endoplasmic Reticulum Tethering Following OPA1 Deletion in Skeletal Muscle. J Cell Physiol 239, e31204 (2024).

56. Huang, H. et al. Cellular succinate metabolism and signaling in inflammation: implications for therapeutic intervention. Front Immunol 15, 1404441 (2024).

57. Mills, E. L. et al. Succinate Dehydrogenase Supports Metabolic Repurposing of Mitochondria to Drive Inflammatory Macrophages. Cell 167, 457–470.e13 (2016).

58. Gobelli, D. et al. The mitochondrial succinate dehydrogenase complex controls the STAT3-IL-10 pathway in inflammatory macrophages. iScience 26, 107473 (2023).

59. Paradies, G., Paradies, V., Ruggiero, F. M. & Petrosillo, G. Role of Cardiolipin in Mitochondrial Function and Dynamics in Health and Disease: Molecular and Pharmacological Aspects. Cells 8, 728 (2019).

60. Jornayvaz, F. R. & Shulman, G. I. Regulation of mitochondrial biogenesis. Essays Biochem 47, 10.1042/bse0470069 (2010).

61. Lai, L. et al. A Role for Peroxisome Proliferator-activated Receptor γ Coactivator 1 (PGC-1) in the Regulation of Cardiac Mitochondrial Phospholipid Biosynthesis* ♦. Journal of Biological Chemistry 289, 2250–2259 (2014).

62. Zhou, S. et al. PGC-1α repression dysregulates lipid metabolism and induces lipid droplet accumulation in the retinal pigment epithelium. Cell Death Dis 15, 385 (2024).

63. Huang, S. et al. Phosphatidate phosphatase Lipin1 alters mitochondria-associated endoplasmic reticulum membranes (MAMs) homeostasis: effects which contribute to the development of diabetic encephalopathy. J Neuroinffammation 22, 111 (2025).

64. Fuentes, J. M. & Morcillo, P. The Role of Cardiolipin in Mitochondrial Function and Neurodegenerative Diseases. Cells 13, (2024).

65. Potting, C. et al. TRIAP1/PRELI Complexes Prevent Apoptosis by Mediating Intramitochondrial Transport of Phosphatidic Acid. Cell Metabolism 18, 287–295 (2013).

66. Schlame, M. & Greenberg, M. L. Biosynthesis, remodeling and turnover of mitochondrial cardiolipin. Biochim Biophys Acta 1862, 3–7 (2017).

67. Regina, A. et al. Cardiolipin preserves Treg metabolic fitness and immune homeostasis in the gut. Nat Metab 8, 1368–1389 (2026).

68. Zhao, W.-Y. et al. Mitochondria-Targeted Antioxidant Peptide SS31 Prevents Hypoxia/Reoxygenation-Induced Apoptosis by Down-Regulating p66Shc in Renal Tubular Epithelial Cells. Cellular Physiology and Biochemistry 32, 591–600 (2013).

69. Jia, Y.-L. et al. The mitochondria-targeted small molecule SS31 delays progression of behavioral deficits by attenuating β-amyloid plaque formation and mitochondrial/synaptic deterioration in APP/PS1 mice. Biochemical and Biophysical Research Communications 658, 36–43 (2023).

70. Schweitzer, G. G. et al. Loss of lipin 1–mediated phosphatidic acid phosphohydrolase activity in muscle leads to skeletal myopathy in mice. FASEB J 33, 652–667 (2019).

71. Evavold, C. L. et al. Control of gasdermin D oligomerization and pyroptosis by the Ragulator-Rag-mTORC1 pathway. Cell 184, 4495–4511.e19 (2021).

72. Hayat. Principles & Techniques of Electron Microscopy. (CRC-Press, 1989).

73. Ngo, J. et al. Isolation and functional analysis of peridroplet mitochondria from murine brown adipose tissue. STAR Protocols 2, 100243 (2021).

