## Supplemental Figures for "Lipin-1-cardiolipin axis sustains mitochondrial respiration and efferocytosis in macrophages"

### SUPPLEMENTAL FIGURE 1

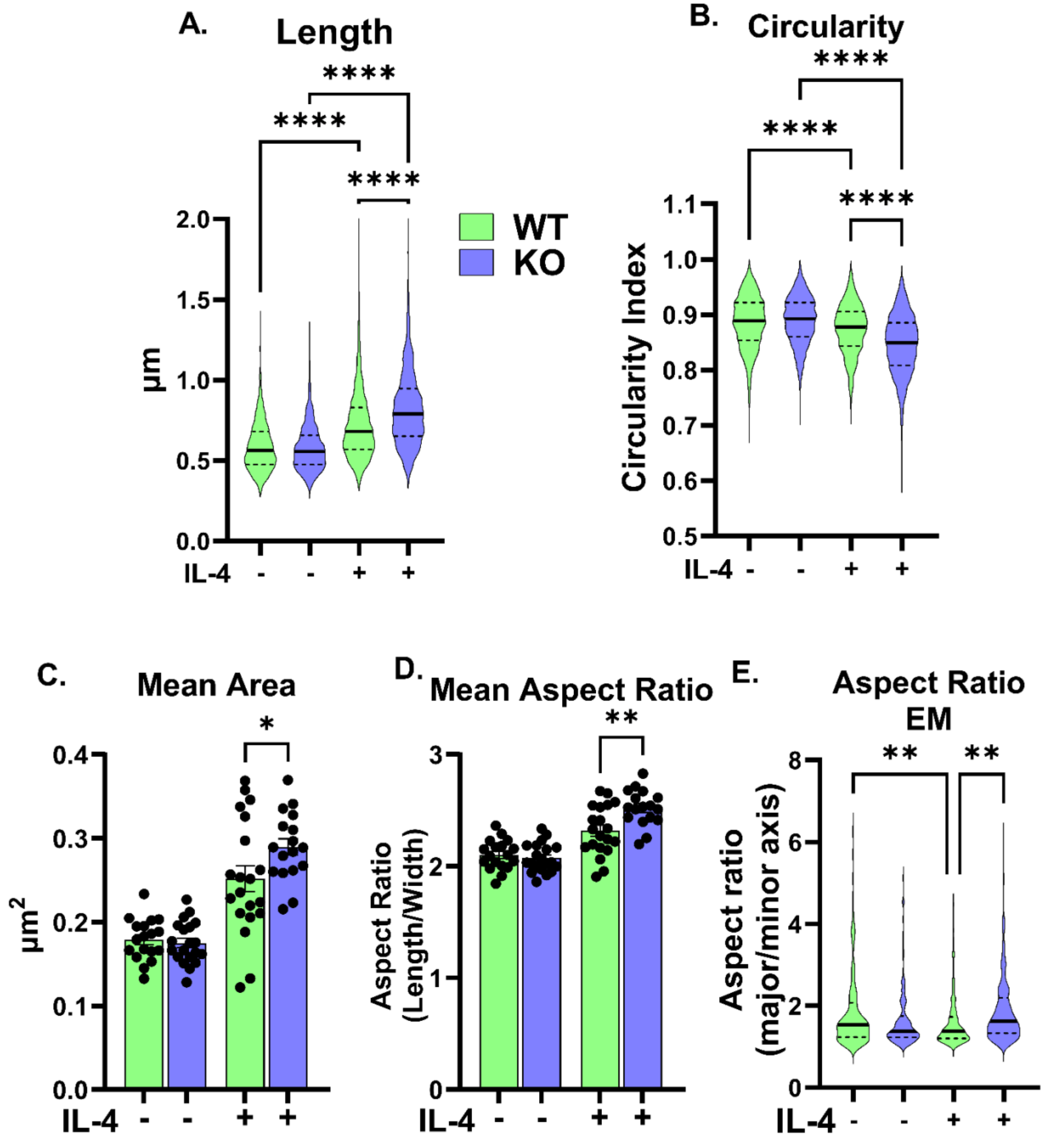

**Supplemental Figure 1. Lipin-1 deficiency is associated with an elongated mitochondrial morphology**

WT and lipin-1mKO BMDMs were treated with or without IL-4 (40 ng/mL) for 6 h followed by confocal imaging of mitochondria identified by TOM-20 immunofluorescence. Violin plots showing the quantification of (A) individual mitochondrial length and (B) circularity. Quantification of (C) mean mitochondrial area and (D) mean aspect ratio. (E) quantification of mean aspect ratio of mitochondria from transmission electron micrographs of untreated and IL-4-treated WT and lipin-1 KO BMDMs. Statistical analyses used are one-way ANOVA. $^*p < 0.05$, $^{**}p < 0.01$, $^{***}p < 0.001$, $^{****}p < 0.0001$; ns, not significant.

**SUPPLEMENTAL FIGURE 2**

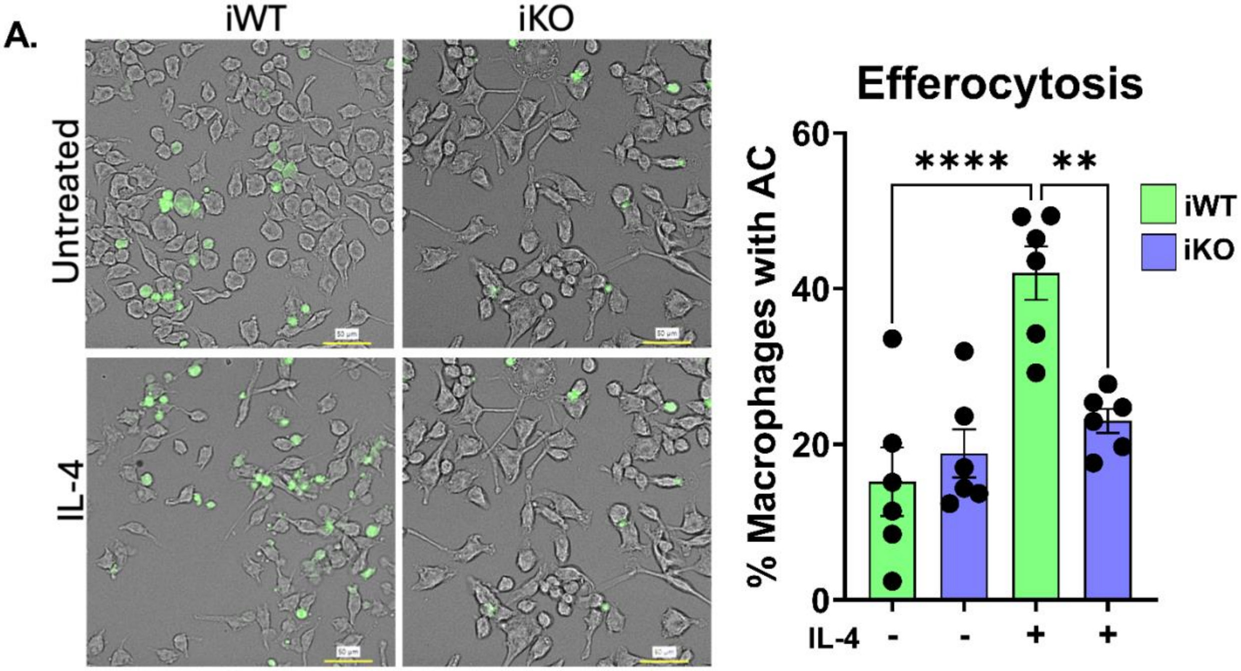


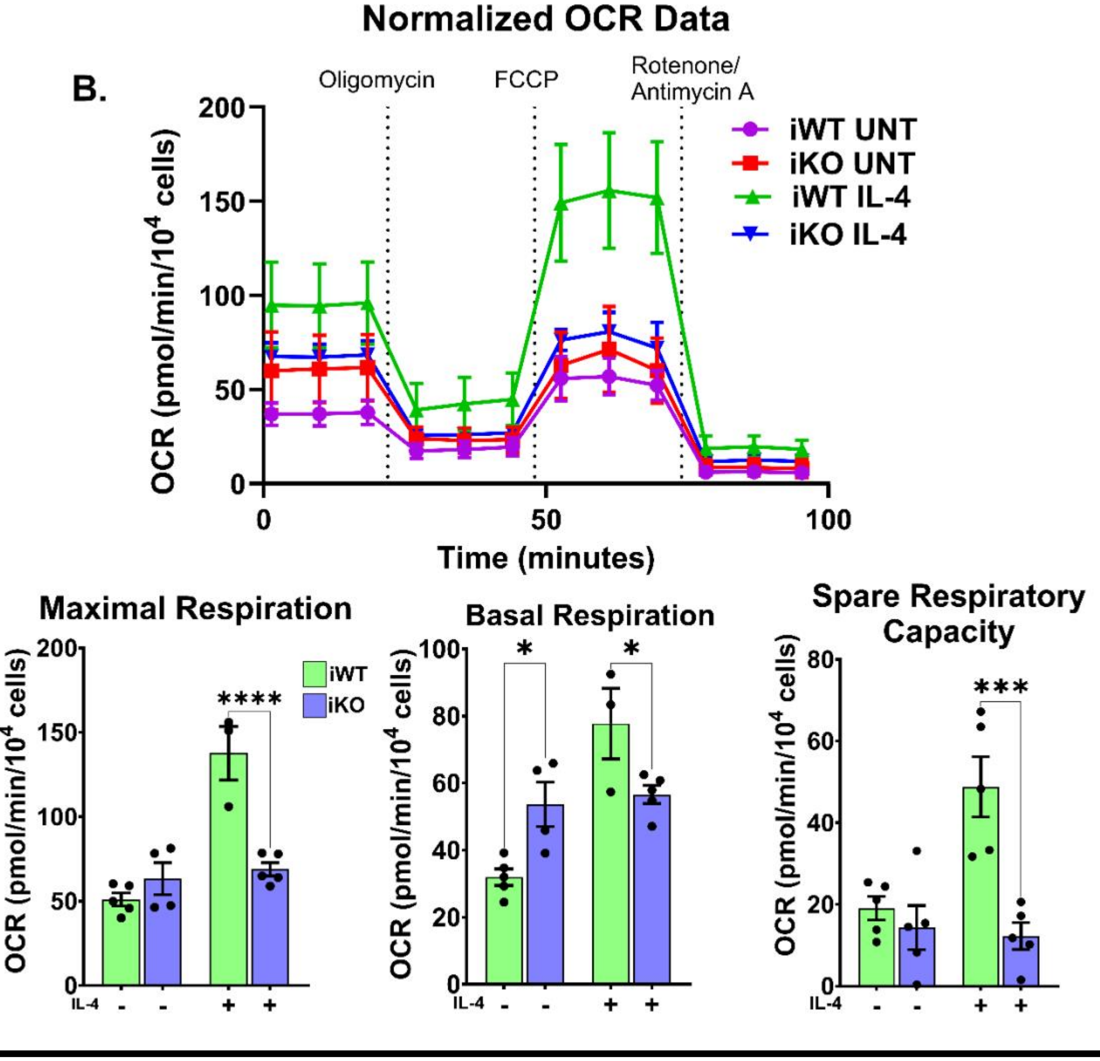

**Supplemental Figure 2. Immortalized WT and Lipin-1mKO BMDMs phenocopy primary WT and Lipin-1mKO BMDMs**

Immortalized WT (iWT) and Lipin-1mKO (iKO) BMDMs were treated with or without IL-4 (40 ng/mL) for 6 h. (A) Representative images of efferocytosis by untreated and IL-4-treated iWT and iKO. CFSE-labeled apoptotic cells are shown in green. Scale bar, 50 µm. Efferocytosis was quantified as the percentage of macrophages containing at least one apoptotic cell, n=6. (B) Mitochondrial respiration was measured by Seahorse XF Mito Stress Test following IL-4 stimulation. OCR was measured during sequential addition of oligomycin, FCCP, and rotenone plus antimycin A. Basal respiration, maximal respiration, and spare respiratory capacity are graphed as means ± SEM. Statistical analyses were done using one-way ANOVA. *$p < 0.05$, **$p < 0.01$, ***$p < 0.001$, ****$p < 0.0001$; ns, not significant.

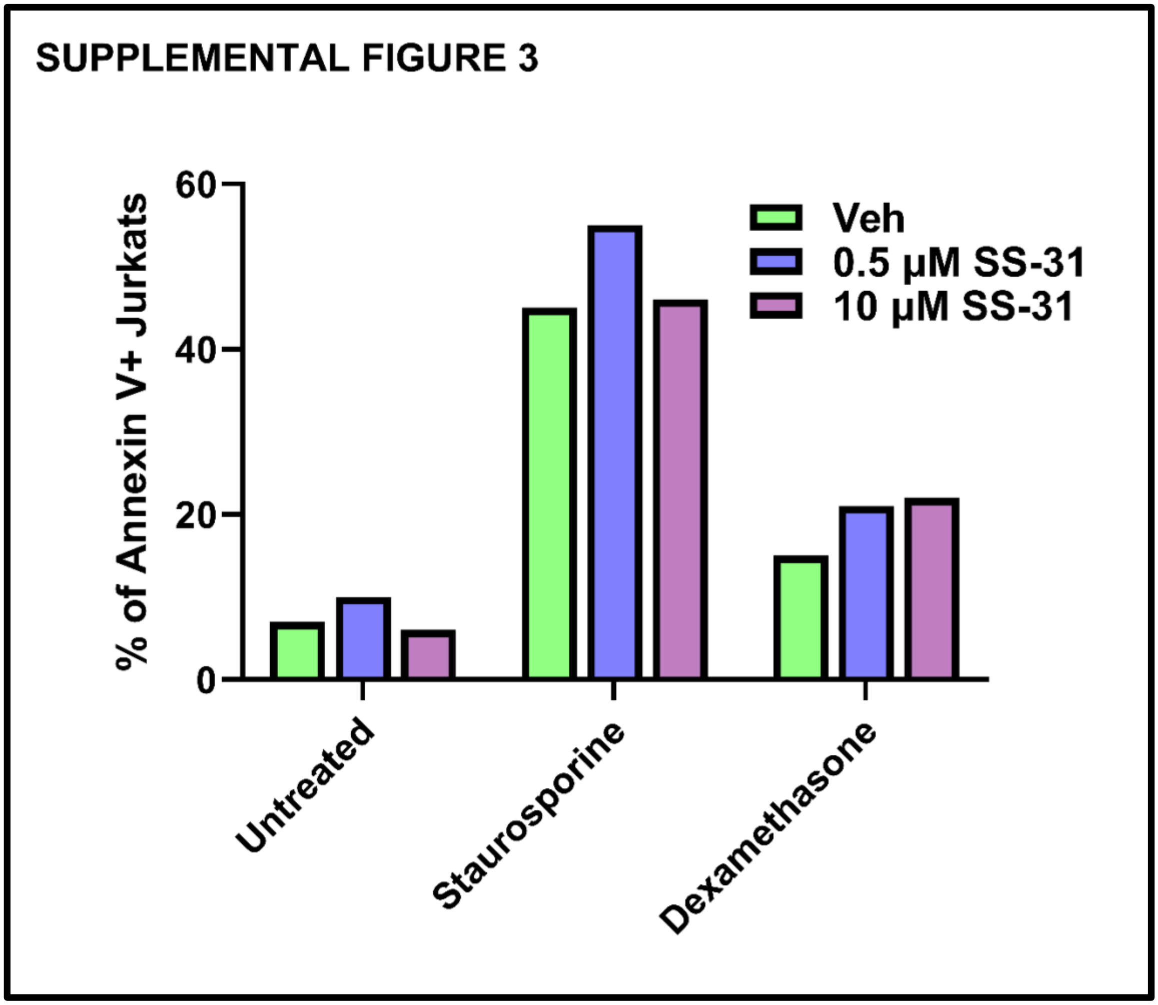


**Supplemental Figure 3. SS-31 does not interfere with staurosporine- and dexamethasone-induced apoptosis**

Jurkats were treated with 0.5 µM or 10 µM SS-31 or vehicle for 2 hours before treatment with 2.5 µM staurosporine, 500 µM dexamethasone, or no treatment for additional 4 hours to induce apoptosis. Apoptosis was measured by Annexin V-binding assay. Graph shows the percentage of Annexin V+ Jurkats
